# Perturbations of a causal synaptic molecular network in autism and schizophrenia revealed with multiplexed imaging

**DOI:** 10.1101/2022.07.11.499330

**Authors:** Reuven Falkovich, Eric W. Danielson, Karen Perez de-Arce, Eike C. Wamhoff, Jeffrey Cottrell, Morgan Sheng, Mark Bathe

## Abstract

The complex functions of neuronal synapses in the central nervous system depend on their tightly interacting, compartmentalized molecular network of hundreds of proteins spanning the pre- and post-synaptic sites. This biochemical system is implicated in the pathogenesis of autism spectrum disorders and schizophrenia, with identified common synaptopathologies and numerous risk genes associated with synaptic function. However, it remains unclear how the synaptic molecular network is altered in these disorders, and whether effects are common to distinct genetic perturbations. Here, we applied PRISM, a quantitative single-synapse multiplexed imaging technique, to systematically probe the effects of RNAi knockdown of 16 autism- and schizophrenia-associated genes on the simultaneous distribution of 10 synaptic proteins. This enabled the identification of novel phenotypes in synapse compositions and distributions. We applied Bayesian network inference to construct and validate a predictive model of causal hierarchical dependencies among eight proteins of the excitatory synapse. The resulting conditional dependence relationships could only be accessed via measurement which is both single-synapse and multiprotein, unique to PRISM. Finally, we show that central features of the network are similarly affected across distinct gene knockdowns. These results offer insight into the convergent molecular etiology of these debilitating, hereditary and highly polygenic disorders, as well as offering a novel, general framework for probing subcellular molecular networks.

## INTRODUCTION

The functional complexity of the brain is enabled by trillions of chemical synapses that form connections amongst its ∼10^11^ neurons. Each synapse is capable of analog computation that integrates its activity history, chemical environment, and the state of its pre- and post-synaptic neurons to modulate communication, for example in classical long-term potentiation (LTP) or depression (LTD). This computation is achieved^1^ in large part by the synaptic molecular network, a dynamic, compartmentalized biomolecular system of hundreds of proteins^2–4^ that includes constantly varying levels and activity states of receptors, scaffolding proteins, kinases, and other protein types. This proteomic diversity likely underlies the remarkable cell- and context-specific functional diversity even amongst synapses of the same type^5,6^. Numerous studies have revealed mechanistic connections between two or three synaptic components at a time, providing the foundation to integrate these connections into a broader context of many-component networks. However, due to the high complexity and interconnectedness of such networks, this integration requires simultaneous single-synapse measurement of numerous proteins, which we previously developed and applied to analyze synapse compositions^7^.

The synaptic molecular network is tightly connected to cognitive disease, with synaptogenesis and plasticity increasingly appreciated as molecular targets for psychiatric treatments^8–10^. Accumulating evidence also points to synaptic biochemistry as a focal point of the pathophysiology of psychiatric, neurodevelopmental and neurodegenerative diseases^11–17^. Autism spectrum disorder (ASD) and schizophrenia (SCZ) are two such conditions that manifest in a range of specific higher cognitive symptoms that range in intensity from healthy neurodiversity to debilitating brain dysfunction. These latter conditions typically include changes in social and communication behavior^18,19^, perception and sensory habituation including self-stimulatory behavior^19–22^, adherence to patterns and focused interests^19^, language acquisition and use^18,23^, as well as general intellectual disability^24,25^ and psychosis^26^. While divergent in symptom presentation, they are often studied genetically in the same context due to similarities in risk genes and possible functional and pathological associations^27–29^.

Both ASD and SCZ are highly heritable^30,31^ and genetically heterogenous^32–34^, with many identified risk genes, including rare, highly penetrant de-novo mutations^35^ as well as many common mutations which contribute small increases in risk. Thus, a central question is whether these different ASD or SSD-causing mutations share similar downstream molecular etiologies, and if so, what are their mechanisms. Genome-wide association studies and rare variant sequencing studies reveal the prevalence of synaptic genes^12,17,29,36–42^, including adhesion, scaffolding, ion channel and local translation control proteins, as well as transcription factors upstream of them, among those associated with ASD and SSD ^43–47^. Additional evidence points to consistent changes in synaptic structural and functional features including dendritic spine morphology^48^, excitation/inhibition ratios^49–52^, and global features of gene expression and protein interaction networks^17,37,53,54^, which are common to different genetic models. Research that implicates perturbations in brain-wide connectivity patterns^55,56^, possibly related to deficits in predictive processing capacity^57^, supports the notion that a fundamental synaptopathology expressed variably throughout the brain may contribute to these disorders^13^. Physical protein interaction approaches such as Y2H and CoIP^53,54^ have established interaction networks involving synaptic proteins and their changes *in vivo* in autistic individuals and autism models, demonstrating the promise of studying the synaptic molecular network as a focal point of autism pathogenesis. However, these studies fall short of measuring changes to joint protein distributions or identifying perturbed causal connections between relevant proteins.

To characterize how synaptic molecular composition is affected across genetic perturbations associated with ASD and SSD, we applied RNAi-mediated knockdown of 16 canonical, highly penetrant risk genes associated with either ASD, SSD, or both, at the onset of synaptogenesis. Once a mature and stable synapse population has been established in each genetic context, we measured the amounts of each of 10 synaptic proteins across individual synapses using Probe-based Imaging for Sequential Multiplexing (PRISM)^7,58,59^. PRISM is a recently introduced multiplexed imaging technique that uses single-stranded DNA (ssDNA)-conjugated antibodies or peptides against desired targets, which are confocally imaged sequentially using fluorescently labeled single-stranded locked nucleic acid (ssLNA) imaging probes. The affinity of ssLNA imaging probes for their complementary ssDNA oligos is ionic strength dependent, allowing sequential rounds of imaging of multiple proteins in the same sample and fields of view by exchanging imaging strands using high- and low-salt buffers. Thus, this imaging method provides a unique combination of extensive multiplexing, moderate throughput, minimal disruption to delicate synapse structures, and single-synapse resolution.

Imaging output consists of images of the same synaptic puncta over numerous protein channels. Integrating PRISM fluorescence intensity over individual puncta and assigning puncta across channels to the same synapse yields individual protein measurements per synapse. The total integrated fluorescence intensity per protein at a given synapse we refer to as the local synaptic protein level. With this approach, in a single experiment we were able to generate a type-resolved systematic view of protein level changes caused by different genetic knockdowns, as well changes to distinct, compositionally defined synaptic populations. These included a global synaptic protein increase with knockdown of *Pten*, a potentially compensatory increase in synaptic PSD95 with knockdown of *Grin2a*, and several unique synaptic phenotypes resulting from *Dyrk1a* knockdown.

Leveraging the unique ability of single-synapse, multiprotein measurements to provide a high-dimensional joint probability distribution (PD) of synapses in composition space, we sought to infer the synaptic protein influence networks that generated the measured protein distributions. To that end we used Bayesian network (BN) inference, a tool previously used to reconstruct entire signaling pathways from multiplexed single-cell data^60^. BNs is a framework to factor a joint PD into a product of individual conditional distributions. This can be represented by a directed acyclic graph between measured nodes, in which graph edges represent direct conditional dependencies between individual nodes, i.e., retaining only those connections which cannot be explained by mutual dependence on a third node, as well as the estimated causal direction of the pairwise dependencies. In this analysis, each node refers to local synaptic levels of a certain protein. Substructures in the resulting model generate testable predictions of causal connections (i.e., the hierarchy in which perturbing A affects B and C) between protein levels, some of which were validated by known protein roles and interactions. In particular, the causal chain in which F-actin determines PSD95, which in turn determines Shank3, we validated independently via direct perturbations, thereby establishing a new causal rule that shapes synaptic protein distributions, generating confidence in the model as a whole. Finally, we present evidence for convergent changes in the inferred synaptic molecular network that are caused by distinct genetic knockdowns, specifically in the strengths of trans-synaptic and intra-postsynaptic edges, offering evidence for a convergent molecular etiology across genes.

## RESULTS

### Effects of ASD- and SSD-associated gene knockdowns on the synaptic molecular system

The following core synaptic proteins were characterized using PRISM to provide snapshots of the synaptic molecular sub-network (**Figure 1A**). Synapsin1 was used to define all synapses, with vGluT1 and vGAT used to differentiate glutamatergic from GABAergic synapses. Other proteins included Bassoon, a central presynaptic scaffolding protein which served as a proxy for active zone size^61^, and the AMPA receptor subunit GluR2, which served as an indicator of synapse strength^62^. Filamentous β-actin (F-actin), measured via phalloidin, was included as the core of the dendritic spine cytoskeleton that is locally regulated by several ASD/SCZ associated genes (e.g., *Trio*, *Pten* and *Dyrk1a*) and whose dysregulation is implicated in various synaptopathologies^63,64^. And, finally, four scaffolding proteins were included that have crucial roles in shaping the post-synaptic density: PSD95, Homer1a, and SHANK3 in excitatory synapses, and Gephyrin in inhibitory synapses. MAP2 staining via conventional immunofluorescence was also used to trace dendrites and constrain puncta assignments to synapses, as well as to align images from different probe exchange rounds^7,58^. For genetic perturbations, ten autism genes (**Figure 1B**) were chosen as the best-scoring targets in the Simons Foundation Autism Research Initiative database^65,66^ (“SFARI score”), half of which are implicated in schizophrenia as well. Six additional schizophrenia-specific genes were chosen from the highly penetrant de-novo mutations identified by Singh et al.^35^.A mixture of four siRNA reagents was used for each gene, and each siRNA treatment was separately validated by RTqPCR for reduction in mRNA levels in cultured hippocampal neurons (**figure S1**). A non-targeting siRNA mix (“NonT”) was also included as a negative control and used for treatment comparisons throughout results.

**Figure 1:**
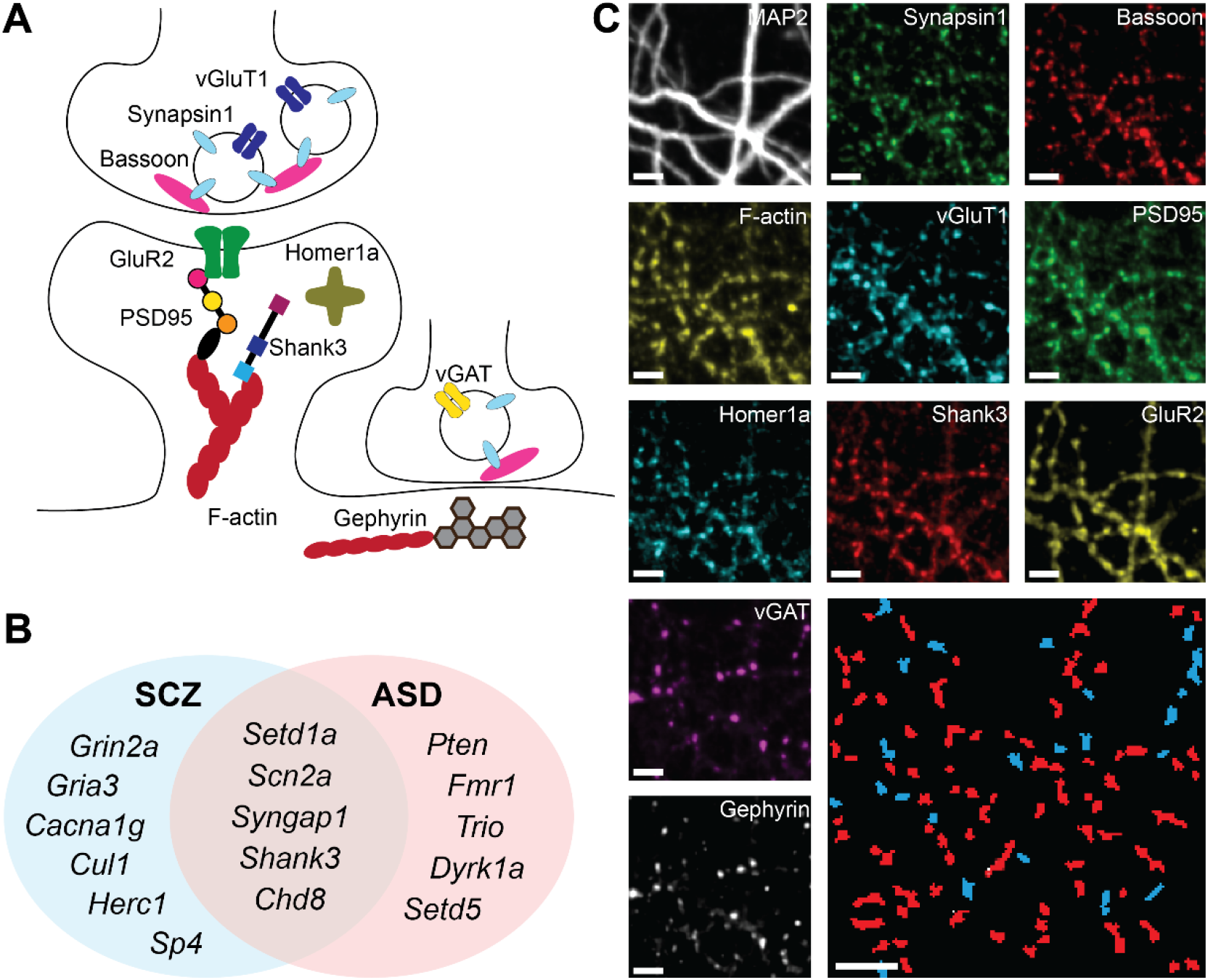
A) Schematic summarizing synaptic roles of the 10 imaged targets. B) Venn agram of gene knockdowns. C) Representative images of the same neuronal culture in ferent imaging rounds, showing colocalized puncta of each protein. Bottom right – automatically identified and segmented excitatory (red) and inhibitory (blue) synapses cale bars are 5µm.

To measure the effect of each gene knockdown on synaptic protein distribution we treated hippocampal neuronal cultures at day-in-vitro (DIV) 6 with one of the corresponding siRNA reagent mixes and fixed the cultures at DIV 19 to image using PRISM (see Methods). We integrated an automated liquid handling platform for probe exchange to complete seven imaging rounds of the same 60 cultures (3-4 per treatment group) in under 12 hours. The resulting fluorescent images of the same synapses across imaging rounds (**figure 1C**) were automatically segmented, classified, and quantified using CellProfiler^67^ (see Methods). We combined data from N=4 such experiments, where each used E18 embryonic neurons from a different pregnant rat, for a total of 3.5×10^6^ 11-protein synaptic measurements across 220 separate neuronal cultures in 18 treatment groups. To account for variability in staining and imaging conditions between experiments, images in each experiment were manually adjusted to the average intensity of that channel in untreated cultures of the same plate.

Automatically segmented puncta, identified as above-threshold intensity peaks within a size range (see Methods) in each protein channel were assigned to specific individual synapses based on overlap with Synapsin1 puncta (see Methods). The existence of a punctum in a protein channel assigned to a specific synapse was taken as the **presence of that protein** in the synapse, and the *integral of fluorescence intensity* of a certain protein channel across its synapse-associated punctum was assumed to reflect the total **level of that protein** in the synapse.

Based on these data, we first examined the individual effects of different siRNA treatments on three global parameters—excitatory:inhibitory (E:I) synapse ratio (**figure 2B**), fraction of GluR2-negative excitatory synapses, implied to be glutamate silent^68^ (2C), and dendrite growth, measured as overall area stained by MAP2 (2D). E:I synapse ratio, implicated previously to be dysregulated in ASD and SSD^49^, was calculated as the ratio of (+vGluT1, -vGAT) to (-vGluT1,+vGAT) synaptic puncta. This ratio averaged ∼5:1 and was significantly increased in knockdown of *Dyrk1a*, consistent with reports of inverse correlation of *Dyrk1a* expression to E:I ratio *in vivo*^50^. It also increased in knockdown of *Grin2a*, *Shank3* and *Chd8*.

**Figure 2:**
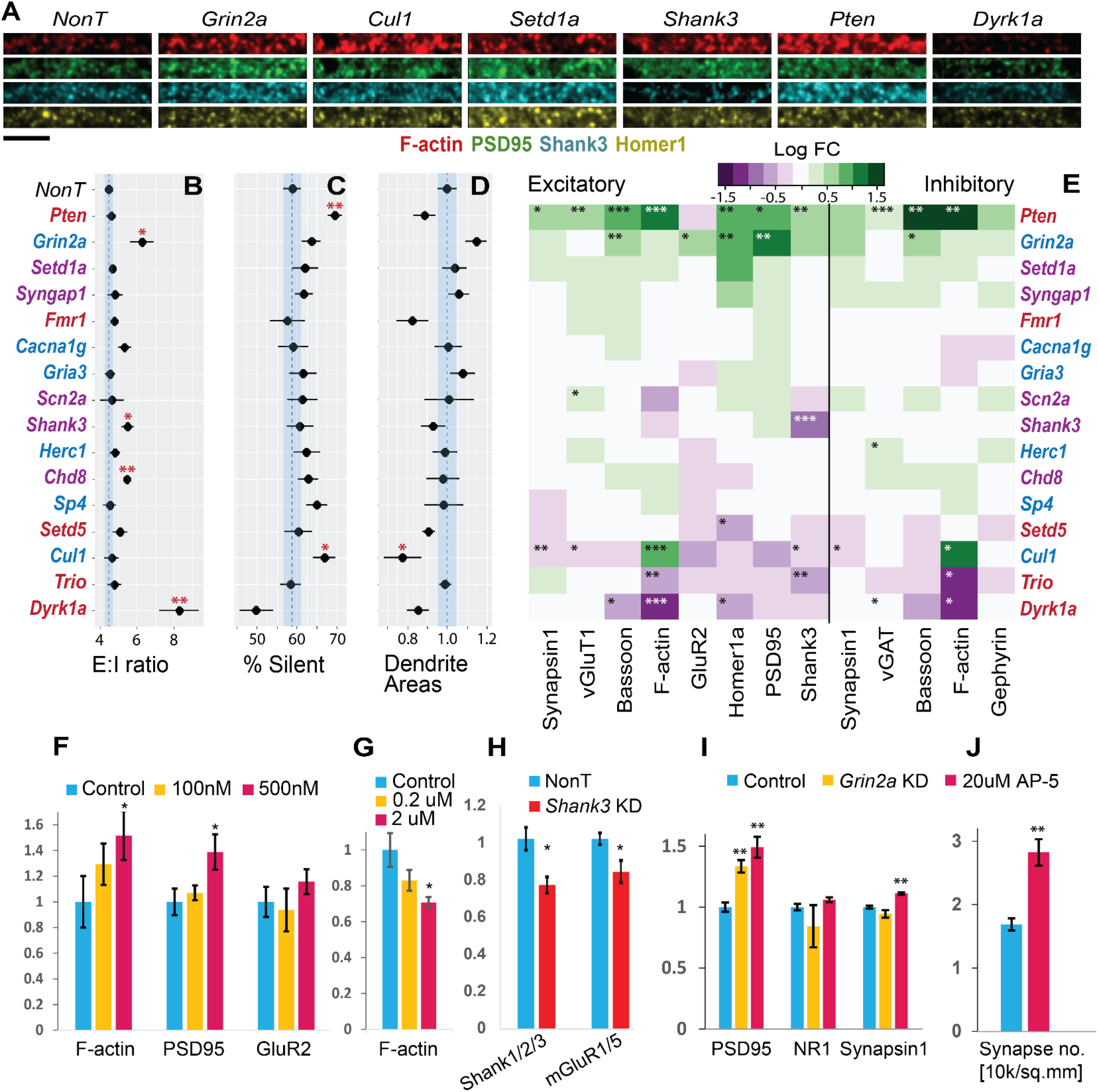
Synapse effects of siRNA knockdown of the genes from figure 1A. **A**) Representative images from medium-sized dendrites across 4 channels. **B**) Excitatory (Syn+, vGlut+, vGAT-) to inhibitory (Syn+, vGlut-, vGAT+) ratios. **C**) Percent of silent (GluR2-) synapses from all excitatory synapses. **D**) Estimates of total dendrite proliferation from MAP2 staining, normalized to NonT. B-D blue lines indicate mean and SEM of NonT measurements. **E**) Log fold change (relative to NonT) of mean levels of each protein in excitatory and inhibitory synapses. **F-J**) Validation experiments with chronic treatments (DIV6-19) by conventional IF. **F-I** average levels of indicated protein per synapse as defined above, in treated cultures normalized to controls. **F**) Three proteins after treatment with 100nM or 500nM bpV(pic), a PTEN inhibitor. **G**) F-actin after treatment with 0.2uM and 2uM Harmine, a Dyrk1a inhibitor. **H**) Shank and mGluR after knockdown of *Shank3*. **I**) PSD95, NR1, and Synapsin1, relative to control after knockdown of *Grin2a* or chronic NMDAR blockade with 20μM AP5. **J**) Density of NR1-positive synapses (in 10^4^ synapses/mm^2^) after treatment with 20uM D-AP5. Error bars indicate SEM across cultures. *p<0.05, **p<0.01, ***p<0.0

Next, by examining the average levels of each measured protein across synapse populations, we created a map of how the synaptic levels of each protein were individually affected by each treatment (**figure 2E**). Beyond the overall characterization, we observed several novel synaptic phenotypes, the strongest of which included: **(a)** a two-fold increase in Homer1a under knockdown of *Setd1a*, a nuclear regulatory lysine methyltransferase, while knockdown of *Setd5*, a gene of the same family, decreased Homer1a. **(b)** An 80% increase in synaptic β-actin after knockdown of *Cul1*, which codes for a component of the E3 ubiquitin ligase complex that has not previously connected to any synaptic protein. Other proteins including Synapsin1 are decreased. **(c)** Knockdown of *Grin2a* led to a ∼70% increase in PSD95, as well as increases in other proteins including Homer1a, GluR2 and Bassoon. **(d)** A twofold decrease in synaptic β-actin following knockdown of *Dyrk1a*, accompanied by decreases in other proteins including Bassoon and Homer1a. **(e)** Knockdown of Trio led to a decrease in synaptic F-actin and SHANK3. We did not identify strongly differential effects on the same protein in the context of excitatory versus inhibitory synapses.

Some treatment effects in **figure 2E** were reported previously or were expected based on pre-existing knowledge of mechanisms involved. For example, we confirmed that siRNA knockdown of *Shank3* led to a marked reduction in SHANK3 levels at synapses. Knockdown of *Pten* led to a broad increase in nearly all synaptic markers, consistent with its role as a negative regulator of PI3K-dependent neurite and synapse proliferation^69^.

To support the validity of PRISM-based phenotyping, we performed knockdown and chemical inhibition experiments using conventional immunofluorescence, showing that: (i) treatment with bpV(pic), a PTEN phosphatase inhibitor, increased synaptic F-actin and PSD95 but not GluR2, mimicking *Pten* knockdown in a dose-dependent manner (2F); (ii) treatment with Harmine, a Dyrk1a inhibitor, decreased F-actin, mimicking *Dyrk1a* knockdown in a dose-dependent manner (2G); (iii) knockdown of *Shank3* reduced synaptic SHANK as well as mGluR1/5, as previously reported^70^ (2H); (iv) knockdown of *Grin2a* increased PSD95 levels, while NR1 levels (corresponding to total NMDAR) and Synapsin1 were not significantly affected (2H). (v) Similar changes were observed with chronic treatment of NMDAR blocker AP-5 (2I), along with a previously reported^71^ increase in density of NR1-positive synapses (2J) that was consistent with our observation of an increased E:I synapse ratio (2B). These last two results suggest a possible compensatory mechanism of PSD95 increase in response to NMDAR deficiency.

### Multiplexed imaging reveals clusters of hierarchical synaptic protein compositions

We applied Uniform Manifold Approximation and Projection (UMAP)^72,73^ on the 11-dimensional dataset of synapse protein levels, yielding the 2D projection in **figure 3A** of different synapse compositions. The distribution of synapses shows distinct clusters defined combinatorially by the presence or absence of certain proteins, similar to our previous observations^58^. These included two inhibitory clusters and several excitatory clusters. Protein absences that defined certain clusters may have resulted from ‘true’ complete absences or merely from levels below threshold. However, we observed similar distributions when changing threshold levels for synapse identification (to 75% and 133% of defined levels) (**figure S2**), indicating that the clusters arose at least in part from qualitatively different synapse compositions.

**Figure 3:**
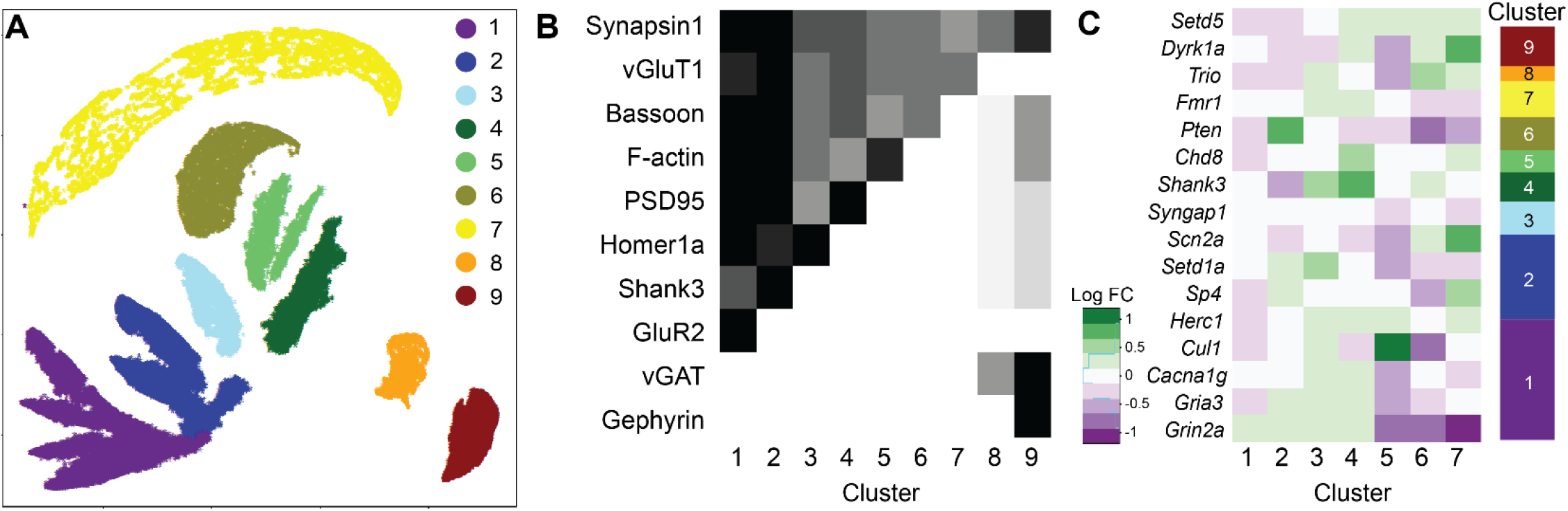
Composition-defined synaptic subtypes. A) UMAP projection of scaled synaptic measurements. B) Relative mean levels of each synaptic protein across each of the clusters identified in A. C) Log fold change of excitatory cluster populations under each treatment. Right – overall composition of the synapse population by cluster (height corresponds to percent of synapses). 01, two-sided t-test.

All treatment groups had synapses in all 9 clusters (**figure S3**). Some population changes between clusters were observed as a result of different gene knockdowns (**figure 3C**). These changes are consistent with the changes observed in mean protein levels in **figure 2E**. For example, *Pten* knockdown, which increased all excitatory proteins save for GluR2, was seen here to enrich cluster #2 (GluR2-negative, positive for all others) at the expense of clusters #1 (GluR2-positive) and #6 (actin-negative). Nevertheless, they provided more detailed information about the specific synapse population changes that occurred as a result of gene knockdown.

In addition, we were able to use spatial information to aid in interpreting protein combinations. A small fraction of puncta was identified as vGluT1-positive, vGAT-negative, and Gephyrin-positive (**figure S4A,B**), despite Gephyrin being well established as an inhibitory synaptic protein^74^. However, upon closer examination we observed that Gephyrin-Synapsin puncta distances in that subset were 50% greater than expected (**figure S4C**), leading us to infer that these puncta were probably not associated physically with the other excitatory markers, and to exclude them from future analyses.

Finally, we observed that not every combination of proteins was present (**figure 3B**). For example, synapses that were negative for Bassoon or β-actin typically lacked or had very low levels of other post-synaptic proteins, indicating a hierarchy in protein dependencies on one another, which we sought to characterize systematically using Bayesian network inference.

### Bayesian network inference of the glutamatergic synapse

All excitatory synaptic proteins were positively correlated with one another (**figure S5A**). However, when we examined each pair individually by measuring their correlation controlled via stratification for all other 6 proteins (see Methods), some correlations disappeared (**figure S5B**). This revealed protein pairs that were not directly correlated but only correlated through their inter-dependence on a third protein, which could be either a common effector (A←C→B) or an intermediate (A→C→B). To systematically map which causal connections among excitatory synaptic proteins were direct versus indirect, and to establish the directionality of their inter-dependence, we derived a Bayesian network from the 8-dimensional distribution of protein levels in excitatory synapses (**figure 4A**).

**Figure 4:**
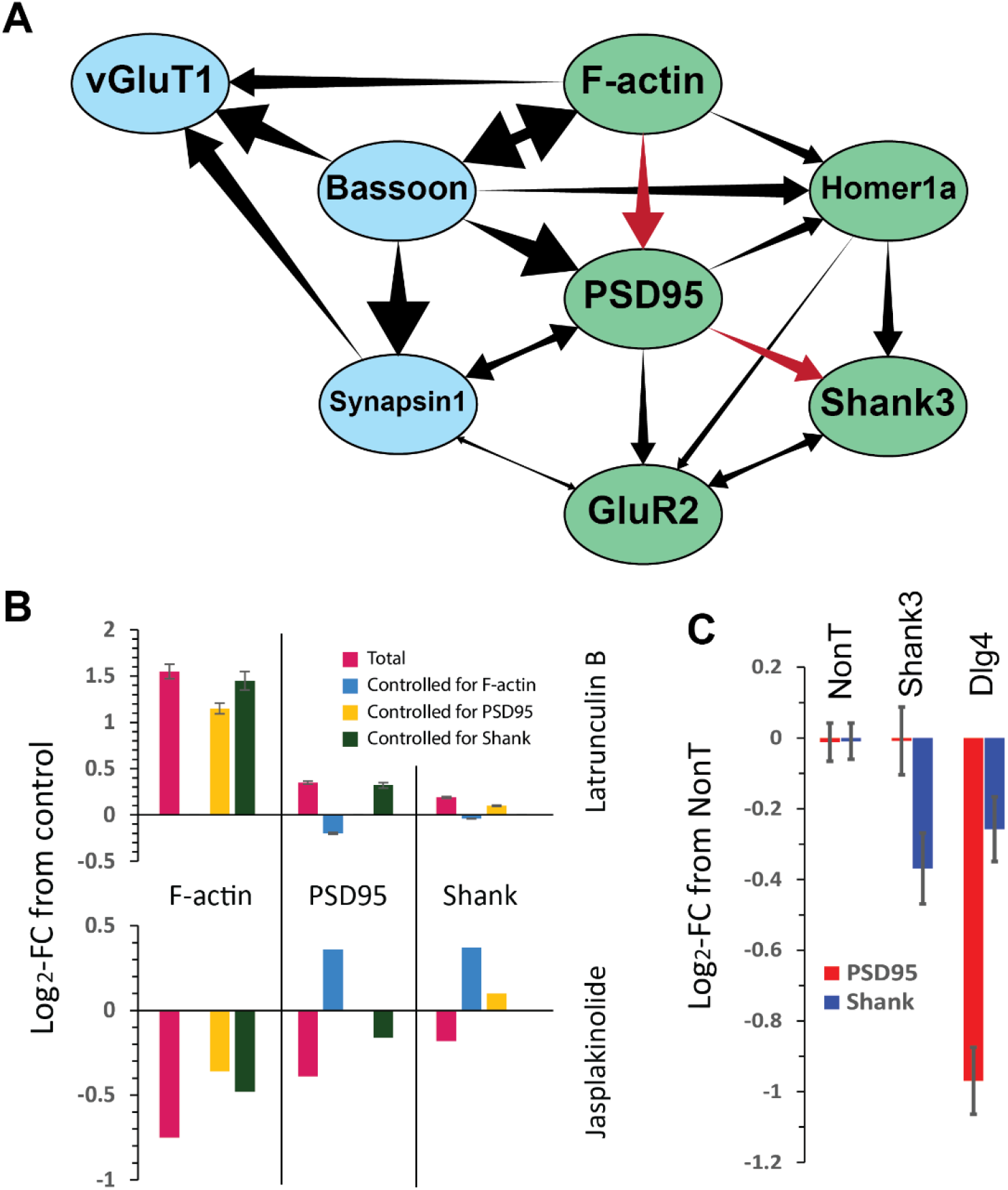
Bayesian network of 8 excitatory synaptic proteins. A) The inferred network. Presynaptic proteins in blue, postsynaptic in green. Red arrows indicate substructure probed in B and C. B) Total and controlled log fold-change in F-actin, PSD95 and Shank3 after treatment with two actin polymerization perturbations. C) Log fold-change in PSD95 and Shank3 after treatment with 3 siRNAs – nontargeting, against Dlg4 (PSD95) and against Shank3. Error bars indicate SEM.

We applied the ‘tabu’ algorithm of the BNLEARN package^75^ in R that searches model-space in a Monte-Carlo-like manner, maximizing an overall score that is based on the likelihood of the data given the model^75–78^. Additional costs to the score were imposed on each edge to prefer simpler, more parsimonious models^75,78^. To estimate confidence levels on the presence of edges and their directions, we applied a bootstrapping method that re-derived separate Bayesian networks for 50 independent samplings of 10,000 synapses from among the 3.5 million measured. An edge was considered present (and is shown in **figure 4A**) if it appeared in >80% of bootstrapped networks, and a direction indicated (as a unidirectional arrow in **figure 4A**) if that direction appeared in >60% of bootstrapped networks where the edge was present. Edge strengths, shown as arrowhead sizes in all network figures, were calculated as the correlations between the parent and child node when controlled by stratification for all other parents of that node^78,79^. In bidirectional edges, both nodes were considered child nodes for this purpose.

This approach was first tested on simulated PRISM-like Bayesian networks—high-dimensional distributions that mirrored real PRISM data but with the distributions of variables conditional on one another in predetermined ways (**figure S6**). A thresholded gamma distribution was chosen for its resemblance to the distribution of the PRISM signal (**figure S6A,B**), and for each ‘simulated synapse’ the value of each variable was sampled from a gamma distribution with a scale parameter determined by values of the corresponding node’s parents. The above method reconstructed predefined Bayesian networks with the right directions (**figure S6C,D**), and edge-strengths calculated as above (with conditional correlations) predicted original interaction parameters (**figure S6E,F**, R^2^=0.92). Notably, even when a distribution was not generated by a Bayesian network (i.e., it contained cycles, which cannot be reflected in a directed acyclic graph but can occur in reality as feedback loops) the inference algorithm still reconstructed the general network structure and edge strengths with reasonable fidelity, reversing some intra-cycle edges to avoid loops but preserving extracyclical edges (**figure S6G,H**).

Network inference on real PRISM data may be sensitive to experimental and image analysis artifacts, such as thresholding, image quality, and rules for synapse identification, as these may impose artifacts on correlations between protein measurements. We therefore performed several quality controls to ensure that our model did not result in such artifacts (**figure S7**). In one, we limited network inference only to synapses positive for all protein components. In another, we varied the thresholds for puncta identification in CellProfiler to 75% or 133% of their values used in the primary analysis. In a third, we used puncta of postsynaptic proteins (F-actin and PSD95) to assign synapse identity, instead of Synapsin1 and vGluT1. All these adversarial manipulations yielded network structures that were largely similar to that in **figure 4A**, especially in presence, relative strengths and directionalities of trans-synaptic and intra-postsynaptic edges. The few inconsistencies included interchanges in the relative positions of Synapsin1 and vGluT1 in the network. Finally, we derived a network based on different 8-protein measurements in a previous study^58^ (**figure S7**). Although that dataset was smaller with fewer perturbations than the current one, reducing confidence in edge presence and directionality, we observed that the six common proteins in both studies exhibited similar connectivity patterns. That the network features were robust against these manipulations, and replicated across experimental conditions, strengthened our confidence that these features represented real underlying biology.

The resulting network (**Figure 4a**) exhibited several features that were anticipated given our knowledge of the function and connectivity of these proteins. For example, the presynaptic proteins Bassoon, Synapsin1 and vGluT1 (the latter two being located together in the same synaptic vesicles) were tightly interconnected, and their correlation was independent of the postsynaptic proteins. Of the three presynaptic proteins, Bassoon levels were most directly ‘causally’ connected to those of the postsynaptic proteins, possibly by acting as a proxy for the size of the active zone. Of the postsynaptic proteins, GluR2 was directly downstream of PSD95, in accordance with the latter’s role as a dynamic anchor for the receptor whose levels dictate the number of sites which can capture diffusing AMPARs on the postsynaptic density^80^. Finally, levels of synaptic F-actin were upstream determinants of all other postsynaptic proteins, possibly due to the cytoskeletal protein acting as a proxy for the size of the dendritic spine which puts a hard limit on other protein amounts. There were also surprising features, such as presynaptic Bassoon seeming to influence levels of postsynaptic F-actin, PSD95, and Homer1 more—from inferred network edge strengths as defined above—than the latter influenced one another.

Most importantly, however, our model lends itself to testable hypotheses of causal connections derived from sub-structures of the network. For example, the position of SHANK3 downstream of the other components predicted that they will not be significantly affected by the direct perturbation of SHANK3, e.g., by siRNA-induced knockdown. Our screen confirmed this (**figure 2F**, *Shank3* line), as does another study that used shRNA knockdown of *Shank3* to show that mGluR5 levels were reduced but other postsynaptic proteins were unaffected^70^. To prevent circular logic (i.e., the presence of the Shank3 knockdown data in the network generating this prediction) we derived the same Bayesian network when excluding the Shank3 treatment group (**figure S9**), showing that BN inference can predict the effects of perturbations that are excluded from the training data.

A more complex predictive sub-structure of the network was the inter-dependence chain: F-actin → PSD95 → SHANK3. This structure predicted that a) perturbing F-actin should affect both PSD95 and SHANK3, but the effect on SHANK3 should decrease when controlling for PSD95, and b) perturbing PSD95 should affect SHANK3, but not vice versa. To test the first prediction, we treated hippocampal cultures with Jasplakinolide (*Jasp.*) or Latrunculin B (*Lat*.) at DIV 6, fixing at DIV 19 and staining for Synapsin1, F-actin, PSD95 and SHANK3 (**figure 4B**). Interestingly, although Jasplakinolide and Latrunculin B are generally inhibitors of F-actin depolymerization and polymerization, respectively, their effects on synaptic F-actin after chronic treatment were reversed. Effects on PSD95 and SHANK3 were consistently in the same direction as the effects on F-actin (decreased for *Jasp*. and increased for *Lat*.) and completely disappeared and even reversed when controlling for F-actin. In addition, the effect on SHANK3 was greatly diminished when controlling for PSD95, as predicted, while the reverse was not true. To test the second prediction, we treated DIV 6 cultures with siRNA mixes against *Shank3*, *Dlg4* (the gene coding for PSD95), or a non-targeting siRNA mix (NonT), and fixed and stained at DIV 21 for MAP2, Synapsin1, SHANK3, and PSD95 (**figure 4C**). Synapsin-controlled SHANK3 levels were reduced compared to NonT in both *Dlg4* and *Shank3*-treated synapses, but PSD95 was reduced only in *Dlg4*-treated synapses, establishing the PSD95>SHANK3 hierarchy. Taken together, these observations supported the conditional dependency chain predicted by the network.

### Convergent effects of siRNA treatments on network structure

Next, we sought to investigate the effects of the siRNA treatments on the protein network informed by its structure. When analyzing the effects of a chemical or genetic perturbation on biomolecular distribution, it is often difficult to distinguish direct from secondary, or downstream effects. Informed by the network model, our multiplexed imaging enables this distinction without requiring additional control experiments. We analyzed the effects of each siRNA treatment on each protein of excitatory synapses while controlling for all parent nodes of that protein in the network, thus retaining only effects that were direct and not mediated by a parent node (**figure 5A**). Isolating direct effects shows us, for example, that *Pten* knockdown affects all the postsynaptic proteins we examined almost exclusively through increased F-actin polymerization in dendritic spines, in accordance with what is known of its regulatory role^69^. It also allows us to discover direct effects which were hidden under second-order effects in the opposite direction, for example that knockdown of *Pten* caused a strong decrease in synaptic GluR2 in parallel to a general increase of all post-synaptic proteins.

**Figure 5.**
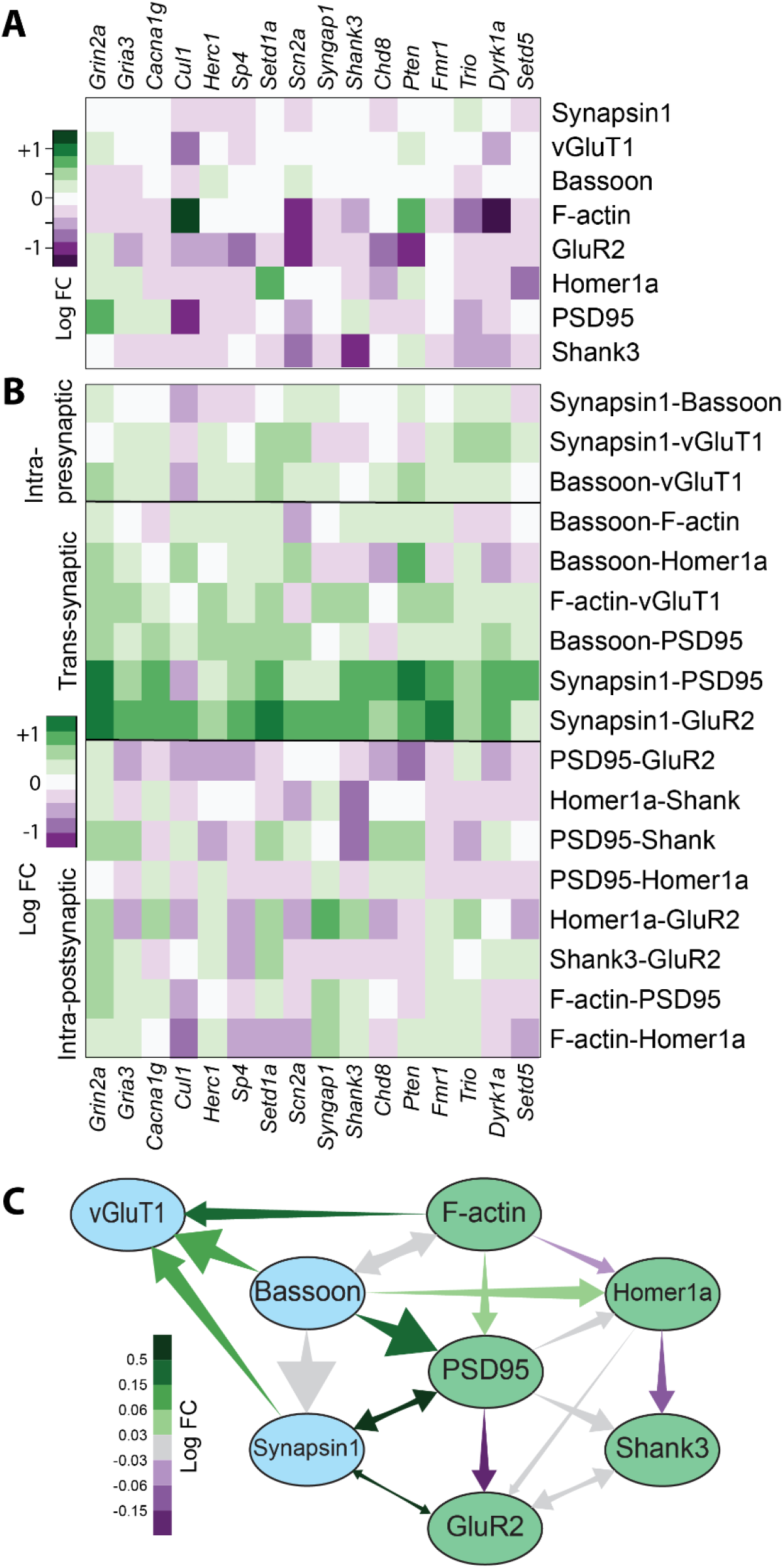
Network-informed analysis of the genetic screen. A) ‘Direct’ effects of each treatment on each protein separately. Similar to 2F but controlled for the parent nodes of each protein. B) Effect of each treatment on the strength of each network edge. C) Network from 4A, each edge colored by the average change in strength across treatments.

Finally, we looked for quantitative effects of siRNA gene knockdowns on the Bayesian network structure itself, particularly on the edge strengths. We quantified the strengths of the 17 edges in the network for each culture (**figure 5B,C**). Surprisingly, we discovered that some edges—particularly centered around PSD95 and GluR2—were uniformly strengthened or weakened by all, or almost all treatments, regardless of direct effects on the proteins themselves. For example, the Synapsin1-PSD95 and Synapsin-GluR2 edges were strengthened in nearly all gene knockdowns compared with the NonT groups, and not weakened in any, even though different treatments increased or decreased each protein on its own. Conversely, the PSD95-Homer1, Homer1-GluR2 or PSD95-GluR2 edges were weakened in most treatments compared with NonT.

Independent treatment effects on different proteins are likely to reduce edge strengths indiscriminately, indicating that that there might be an underlying molecular process by which knockdowns of many different genes all serve to, for example, weaken the extent by which PSD95 determines GluR2 and strengthen the extent by which PSD95 and Synapsin1 influence each other, where these effects are superimposed on any direct effects they may also have on these proteins. To the best of our knowledge, this is the first time that a direct measurement of synaptic protein networks has yielded molecular phenotypes common to many different ASD- and schizophrenia-associated mutations, which may be reflective of shared synaptic pathogenesis.

## DISCUSSION

### Detailed parallel phenotyping of synaptic biochemistry under ASD and SSD models

We used simultaneous measurement of multiple proteins at single-synapse resolution to infer causality relationships among protein numbers in the synaptic molecular network, and systematically map how it is affected by the perturbation of autism- and schizophrenia-associated genes. PRISM, supported by automated probe exchange, image analysis and synapse segmentation and quantification, can be used to measure multiple synapse-specific phenotypes across many treatment groups in a single experiment, as well as identify synapse types and population changes that are more subtle than bulk effects on a protein due to its ability to resolve synapse-to-synapse heterogeneity in multiple protein levels.

We observed phenotypes that were anticipated based on previous studies or their known activity, such as *Pten*^69^ and *Shank3*^70^. We also observed phenotypes that to the best of our knowledge have not been previously reported, such as *Cul1* on F-actin, *Setd1a* on Homer, and *Grin2a* on PSD95. Of the overall molecular signatures, *Dyrk1a* exhibited the most unusual pattern, combining a markedly higher E:I ratio, lower levels of F-actin and downstream postsynaptic proteins, but not of GluR2 (and generally a lower percentage of silent synapses). This is congruent with reports of *Dyrk1a*-associated autism presenting a unique neurological character^81,82^. While the effects from the preliminary screen in our model system of siRNA knockdowns would need to be reproduced in truer *in vivo* genotypes (e.g., many of these are loss-of-function mutations or deletions leading to haploinsufficiency, which may be reasonably modeled by siRNA knockdown. Thus we expect the effects of these on fundamental synaptic biochemistry to be similar to those observed from more representative mutations.

Another important consequence of the ability to measure multiple proteins simultaneously is that it can facilitate deconvolution of direct causal relationships from those mediated by other processes. Instead of having to experimentally constrain any possible confounding variables, multiplexed imaging allows measuring them simultaneously, while moderate throughput offers a large enough number of data points (>3·10^6^ in this study) to directly control for possible confounding variables via stratification. The results of such intra-dataset controls must be interpreted carefully due to potential artifacts arising from comparing different synapse populations. Nevertheless, when using information about causal connections between variables, either from prior knowledge or, as in this case, inferred directly from the probability distributions, such a controlled analysis can help identify when a certain effect is entirely mediated by other variables (as with *Pten* or *Dyrk1a* on postsynaptic proteins, mediated by F-actin) or when a certain variable deviates significantly due to a hidden direct effect from what is expected given its upstream network connections (as with *Pten* on GluR2).

### Interpreting Bayesian network structure and edge strengths

In the inferred Bayesian network, an edge from node A to B indicates that the PD of B depends on A even after accounting for all other measured components. This dependence can be direct or via a hidden (unmeasured) component. Conversely, lack of a direct edge between two correlated nodes indicates that any correlation between them can be explained by the other known components. Thus, while the overall structure is dependent on the set of proteins measured, we expect ‘non-edges’, such as F-actin-SHANK3, to be a conserved feature even as the network expands to include more proteins. The structure we observed hints at some general rules governing synaptic molecular composition, which have also been observed in the literature: relative independence of the presynaptic active zone assembly and molecular composition^83,84^, and receptor levels driven by scaffolding proteins rather than the reverse.^80,85,86^

The interpretation of changing edge strengths is not mechanistically obvious. As a rule, injecting perturbative noise into a system weakens correlations by default. Thus, a weakened edge may correspond to a loss of correlation that can occur if only one protein of a pair is perturbed, or if they are perturbed in different directions. However, this is not always the case—for example, knockdown of *Cul1* reduces both PSD95 and GluR2 to a similar extent, but also weakens the PSD95-GluR2 edge. In general, an edge from a parent node to a child node is considered in the context of all the other parents of that node. Thus, an effect that perturbs the mechanism by which one parent node affects the distribution of the child will weaken the corresponding edge but may strengthen the edge from another parent, and vice-versa. With this in mind, it is interesting to observe that trans-synaptic edges are strengthened by the genetic knockdowns, while intra-postsynaptic edges are generally weakened.

### Comparison with other protein network models

Our results feed into a growing body of genomic and proteomic observations of convergent changes at the protein network level in autism. These are based on physical interaction networks derived from yeast two-hybrid (Y2H) tests^53^ or bulk quantitative multiplex co-immunoprecipitation (QMI)^54^. It is important to note the difference between physical interaction networks as obtained from Y2H screens or coimmunoprecipitation, and our network, which does not provide information about protein interactions, but rather about the constraints on the multiprotein joint PD. When many parallel interaction pathways are known, the Bayesian network allows to select one that causally determines protein levels. For example, the F-actin → PSD95 → SHANK3 causal chain we established implies that it is likely an interaction chain from actin to PSD95 (possibly via ARPC4^87^) that drives PSD95 (and therefore SHANK3) levels. It must be noted that for actin, our measurement mixes together signal from both pre- and post-synaptic β-actin filaments. However, since postsynaptic F-actin is much more abundant, we believe the observed conditional dependencies are due mostly to postsynaptic F-actin levels.

Causal correlations may also appear without known interactions if the measured target is the best available proxy for a different measure that drives protein levels. For example, no direct or even singly-mediated interactions of Bassoon with F-actin, PSD95 or Homer1a are known or expected. Rather, we hypothesize that these edges represent different paths, through unmeasured targets, by which the size of the active zone (for which Bassoon is a proxy) affects levels of different post-synaptic proteins. Finally, our network so far only measured protein abundances, and not phosphorylation activation or other modification changes (or splice isoforms), which are often influenced by interaction partners.

The two frameworks thus complement each other; physical interaction networks provide mechanistic details and, by casting a wider, less biased net, identify new causally important proteins to measure, while Bayesian network analysis integrates these details into a global picture of what shapes the overall synapse protein composition. This, in turn, provides insight into possible dynamic processes in the synaptic molecular network, which are tested by once again returning to mechanistic connections between components.

### The steady state of the synaptic molecular network in ASD and SSD

We propose a theoretical framework for future exploration into the convergence of phenotypes of different disease-associated mutations at the level of the synaptic molecular network. A dynamic system with many different components such as the synapse exists in a very high-dimensional space of component abundances and activation states. However, with sufficient interactions and feedback loops, provided by the high degree of interconnectivity and interactions observed among synaptic proteins, and especially noting feedback constraints imposed by the biology of the synapse (e.g., homeostatic plasticity), such systems invariably settle into a much lower-dimensional space of allowed stable or metastable states. In other words, since synaptic structural dynamics (e.g., LTP and LTD) occur at longer timescales than the biochemical feedback interactions which establish the allowed states, the former navigate a comparatively narrow landscape of only those states which the dynamic system can consistently sustain, and the synapse population distribution reflects this landscape.

A similar constraint induced dimensionality reduction is considered to stabilize symmetric phenotypes in genotype-phenotype maps^88^. In the context of ASD- and schizophrenia-associated genotypes mapping to a synaptic phenotype space with reduced dimensionality caused by interaction-based constraints, this means that a multitude of seemingly unrelated disease-associated mutations, can drive similar perturbations to the lower-dimensional stable/allowed-state landscape, as measured in synaptic protein networks.

Thorough investigation of such a mechanism in autism and schizophrenia will require characterizing this space of allowed synaptic states and how it changes under different mutations, a characterization for which this work and others provide initial outlines. This characterization would also provide additional starting points to investigate the downstream effects of convergent synapse structure phenotypes on the architecture of the synaptome and specific neuronal circuits.

Given the potential of PRISM with automated probe exchange for increasingly higher throughput, along with the theoretically unlimited multiplexing capacity, our method of multiplexed imaging with single-synapse analysis is well-poised to investigate such hypotheses, as well as any other processes in the synaptic molecular network. Specifically, the properties of BN inference mean that additional measured nodes would increase both the scope of, and confidence in, network structures, as well as help establish differences in network structure underlying qualitatively different functionality in excitatory synapses connecting different neuron types^6^. Finally, the tool presented here needs not be limited to synapses, and can be applied to any subcellular structure which can be identified individually in fluorescent microscopy, including mitochondria, phagosomes, nuclear compartments, etc. Thus, PRISM, supported by Bayesian network analysis may come to serve as a fully general hypothesis-generating tool for understanding complex protein networks in situ in cells, organelles, and subcellular structures.

## ACKNOWLEDGEMENTS

This work was supported by NIMH R01 grant MH112694-05 and NSF Physics of Living Systems grant 1707999. The authors would also like to thank Agnes Walsh for assistance in preparing DNA-conjugated antibodies, Juliana Strother for assistance in validation experiments and Douglas Lauffenburger for advice on Bayesian Network inference.

## METHODS AND SUPPLEMENTARY FIGURES

### Methods

#### Antibody-docking strand conjugation – SMCC

Antibodies were purchased as formulations without serum proteins as listed below (Table S1), 0.1-1 mg of antibody were purified into PBS using Zeba spin columns (7 kDa, Thermo Fisher Scientific). Subsequently, antibodies were concentrated to 1 mg/ml using Amicon Ultra centrifugal filters (100 kDa, 4000 g, EMD Millipore). The initial concentration of anti-vGAT was 2 mg/ml. From a freshly prepared stock at 2 mM in DMF, SMCC (Sigma Aldrich) was added to the antibody at 7.5x molar excess. The reaction mixture was protected from light and incubated for 3 h at 4°C on a shaker. Excess SMCC was removed by purification into PBS using Zeba spin columns (7 kDa, Thermo Fisher Scientific). In parallel, 25 nmol 5’ thiol-modified ssDNA (Integrated DNA Technologies, modification catalog no. /5-ThioMC6-D/) was dissolved in 25 ul water and 55 ul PBS with 2 mM EDTA at pH 8.0 (Table S2). After the addition of 20 ul of a freshly prepared stock of 500 mM DTT in PBS with 2 mM EDTA at pH 8.0, the reaction mixture was protected from light and incubated for 2 h at 25°C on shaker. The reduced 5’ thiol-modified ssDNA was purified into water using NAP-5 columns (GE Life Sciences). Fractions containing ssDNA were identified using absorbance measurements at 260 nm and DTT was monitored calorimetrically using bicinchoninic acid. The reduced 5’ thiol-modified ssDNA was immediately added to the antibody-SMCC conjugate at 15x molar excess, the reaction mixture was protected from light and incubated overnight at 4°C on a shaker. Antibody-ssDNA conjugates were purified into PBS using Amicon Ultra centrifugal filters (50 kDa, 4000 g, EMD Millipore). Amino-modified phalloidin (Bachem) was conjugated using the procedure described above, but with the following changes: the molar excess of SMCC was 10x and the molar excess of reduced 5’ thiol-modified ssDNA was 1x. HPLC purification was employed to remove unreacted SMCC and 5’ thiol-modified ssDNA, respectively (Waters, BEH C18 column, gradient for phalloidin-SMCC: from 80% TFA in water and 20% acetonitrile to 20% TFA in water and 80% acetonitrile over 10 min, gradient for phalloidin-ssDNA: from 90% 0.1 M TEAA in water and 10% acetonitrile to 60% 0.1 M TEAA in water and 40% acetonitrile over 10 min). Antibody concentration were determined by absorbance measurements at 280 nm. Conjugation efficiency was estimated by MALDI-TOF mass spectrometry and ranged from 1 to 3, depending on the antibody. Antibody-ssDNA conjugates were stored at −20 C in PBS with 50% glycerol.

#### Antibody-docking strand conjugation - SiteClick

For anti-Homer 1, the ssDNA conjugates using the SiteClick technique (Invitrogen) conjugation method following the manufacturer’s instructions. Briefly, 200 μg of the anti-Homer 1 was concentrated to 2 mg/ml in 1x Tris buffer and incubated with β-galactosidase. Azide-modified, terminal galactosides were attached using β-galactosyltransferase. Azide-modified antibody was purified into 1x Tris buffer using Amicon Ultra centrifugal filters (50 kDa, 4000 g, EMD Millipore). 5’ DBCO-modified ssDNA (Integrated DNA Technologies, modification catalog no. /5-DBCON/) was dissolved in water, added to azide-modified antibody at a molar excess of 30 and incubated overnight at 25°C (Table S2). Antibody-ssDNA conjugates were purified into PBS using Amicon Ultra centrifugal filters (50 kDa, 4000 g, EMD Millipore). Antibody concentration were determined by absorbance measurements at 280 nm. Conjugation efficiency was estimated by MALDI-TOF mass spectrometry and ranged from 1 to 2, depending on the antibody batch. Antibody-ssDNA conjugates were stored at −20°C in PBS with 50% glycerol.

#### Imager strands

25 nmol of 5’/3’ diamino-modified ssLNA (Qiagen) was dissolved in 500 ul PBS with 10% DMSO at pH 8.3 and 250 nmol of NHS-Atto 565 or NHS-Atto 655 (Sigma Aldrich) were added from a 15 mM stock in DMSO (Table S2). Following immediate vortexing, the reaction mixture was protected from light and incubated overnight at 25°C on a shaker. Excess dye was removed using NAP-5 columns (GE Life Sciences). Fractions containing ssDNA were identified using absorbance measurements at 260 nm. Subsequently, 0.1 M TEAA was added ssLNA-dye conjugates and conjugates bearing two dyes were purified by HPLC (Waters, BEH C18 column, gradient for Atto 565: from 80% 0.1 M TEAA in water and 20% acetonitrile to 70% 0.1 M TEAA and 30% acetonitrile over 10 min, gradient for Atto 655: from 90% 0.1 M TEAA in water and 10% acetonitrile to 75% 0.1 M TEAA in water and 25% acetonitrile over 10 min). Peaks corresponding to ssLNA conjugates bearing two, one or no dye were assigned based on absorbance spectra. Solvents were removed in vacuo and ssLNA-dye conjugates were dissolved in water at 10 to 100 µM, depending on the yield. Yields were determined by absorbance measurments at 565 nm or 655 nm.

#### Neuronal culture and treatment

Procedures for rat neuronal culture were reviewed and approved for use by the Broad Institutional Animal Care and Use Committee, in accordance with the National Institutes of Health Guide for the Care and Use of Laboratory Animals. In each of N=4 biological repeats, 1-2 Embryonic Day 18 embryos were collected from a separate pregnant Sprague Dawley rat killed by CO_2_ (Taconic). Embryo hippocampi were dissected in 4°C Hibernate E supplemented with 2% B27 supplements and 100 U/ml penicillin/strep (Thermo Fisher Scientific). Hippocampal tissues were digested in Hibernate E containing 20 U/ml papain, 1 mm L-cysteine, 0.5 mm EDTA (Worthington Biochem), and 0.01% DNase (Sigma-Aldrich) for 8 min. Neurons were centrifugated at 1000rpm by 5min, pellet with cells were then resuspended into NbActiv1 (BrainBits LLC, now TransnetYX) supplemented with 25mM glutamate, and plated at a density of 15,000 cells/well onto poly-d-lysine-coated, black-walled, thin-bottomed 96-well plates (Corning BioCoat). After 48 hours, AraC was added to each culture at a concentration of 1uM, to suppress glia proliferation and minimize well-to-well variability resulting from it. At DIV 5, the media was entirely replaced with warm NbActiv4. At DIV 6, each culture was treated with Accell SMARTpool (Dharmacon/Horizon from Perkin Elmer), a mix of four chemically modified self-transfecting siRNAs, against the relevant gene (Table S3) to a total siRNA concentration of 1uM in NbActiv4. Cultures were then left unperturbed until fixation on DIV 21. Each plate included 60 wells/separate cultures, 3-4 in each treatment group. Across 4 plates, one for each biological repeat, this results in a total of n=11-18 technical repeats in each treatment group.

#### RTqPCR knockdown validation

RTqPCR was performed using Fast Advanced Cells-to-CT kit (Ambion) according to the manufacturer’s protocol. In short, cells were prepared for lysis by washing them with cold PBS 1x, then Stop solution was added following lysis buffer with DNAse I. RT Master Mix using Cells-to-Ct lysate was prepared and reverse transcription was done on a thermal cycler. Lastly, qPCR was done using LightCycler® 480 Probes Master (Roche) with TaqManTM Gene Expression Assays designed for each target (see Table S3 for catalog numbers) and performed on a LightCycler® 480 Instrument. Two TaqManTM Gene Expression Assays (Life Technologies). *Actb* was used as a reference gene to normalize the results (Life Technologies). For relative quantification of gene expression, the 2− ΔΔCt method was used^1^.

#### Staining and Imaging

Cells were fixed and stained as described previously^2–4^. Briefly, cells were fixed in paraformaldehyde with sucrose and permeabilized with Triton X-100. They were then incubated in a mixture of RNases A and T1 to reduce the fluorescent background caused by ssLNA-RNA binding and blocked with 5% Bovine Serum Albumin (BSA). The first round of primary staining was performed using unconjugated primary antibodies (table S1 rows 1-6) diluted in the regular blocking buffer. Cells were blocked with nuclear blocking buffer [5% BSA and 1 mg/ml salmon sperm DNA (Sigma-Aldrich) in PBS] and then incubated with conjugated secondary antibodies (table S1 rows 7-11) diluted in the nuclear blocking buffer. After post-fixation, cells were stained in the third round with conjugated primary antibodies (table S1 rows 12-16) in nuclear blocking buffer and then with DAPI.

High-throughput spinning disk confocal LNA-PRISM imaging was performed using the Opera Phenix High-Content Screening System (PerkinElmer) as described before^4^ with the following main changes: first, two colors were used for PRISM in each round, and second, probe introduction, wash and exchange was performed automatically using a Bravo automated liquid handling system. In each round, a pair of imaging probes in two colors (see table S2 for sequences) was freshly diluted to 10 nm in imaging buffer (500 mm NaCl in PBS, pH 8) immediately before imaging. Neurons were incubated with imaging probes for 5 min and then washed twice with imaging buffer to remove unbound probe. The plates were then imaged in 4 wavelengths: 405nm (DAPI), 488nm (MAP2), 561nm (orange probe) and 647nm (red probe). For each field of view, a stack of five images was acquired with an axial step-size of 1 μm. Either four (in one plate) or nine (in the other three plates) lateral fields of view were imaged in each culture. Following each round of imaging, cells were washed two times with wash buffer (0.01 × PBS) for 3 min per round, and then re-imaged to ensure that all PRISM fluorescent signal was removed before introducing the next probe pair. After all imaging rounds, neurons were stained with a 568nm fluorescent nanobody against vGlut1 (table S1 row 17) for 1 hour and imaged again.

#### Data exclusion

The following subsets of images were excluded from analysis:

- One row in plate #1, for which one of the imaging rounds was out of focus.
- All images of NR2A, for which staining was very diffuse and very few puncta could be identified.
- All images of cultures treated with siRNA against Xpo7, which exhibited highly irregular staining patterns for Homer1 that could not be reproduced with other batches of that siRNA. We attributed the effect to an issue with the specific siRNA batch used.

#### Automated image analysis using cellprofiler

CellProfiler was used to automatically correct, align, segment and quantify synapses in images. This tool allows for modular construction of pipelines for image analysis^5–7^. The pipeline used here is similar to a previous study^4^ and is available on github.com/lcbb. The main steps in the image analysis pipeline are as follows:

1. By-pixel maximum projections of confocal Z-stacks of all images in each round are calculated separately and loaded into CellProfiler.
2. MAP2 (488nm) images in each round are used to align images of other channels between rounds.
3. An illumination profile correction is applied to all images based on background averages across all wells.
4. For each round and wavelength, the average intensity in untreated wells of a plate is calculated and used to normalize the images in all other wells. This is used to account for between-plate differences in exogenous brightness (staining strength, laser strength, exposure time etc.)
5. The DAPI image is used to identify nuclei objects. All other images of the same field are then masked by the nuclei.
6. The MAP2 image is used to identify dendrite objects.
7. A white top hat filter with a radius of 4px is applied to all synaptic protein images across all rounds to enhance puncta.
8. For synapse counting analysis (figure 2B-E), synaptic objects were segmented and identified in images of each channel by applying the RobustBackground tool, which calculates an optimal threshold value for each image individually based on the intensity histogram. For all other analyses, we calculated a per-channel global threshold from the average threshold calculated by RobustBackground across all imaged fields in untreated wells. We then applied this value as a uniform threshold to all images of that channel to ensure that all images are segmented identically.. Only
9. Synapsin1 puncta are then masked using the dendrites previously identified, to retain only puncta which are within 12px of a dendrite. These are then defined as synapses.
10. Puncta in all other channels are assigned to synapses if they overlapped with Synapsin1 puncta more than 6.25% (for postsynaptic proteins) or more than 50% (for presynaptic proteins).
11. Finally, levels of each protein per synapse are calculated as the intensity integral of that protein’s image across its puncta. If a certain protein did not have an identified puncta associated with a synapse, its level was marked as 0.

Synapses were identified as excitatory if they contained only vGlut, inhibitory if they contained only vGAT, and otherwise excluded from further analysis (positive or negative for both vGlut and vGAT). Excluded synapses were 20-30% of all identified synapses. We also performed the same analysis with uniform threshold values of 75% and 133% of the calculated average, which yielded more and less synaptic puncta, respectively, but similar observations in treatment effects, clusters, Bayesian networks and edge strengths. In controls for network inference by alternative synapse identification, instead of Synapsin1 puncta for synapse definition and assignment of all other proteins, we used postsynaptic puncta defined by merging of F-actin, PSD95 and Shank3 puncta.

#### Statistical Analysis

Unless stated otherwise, all statistical comparisons among treatment groups are for groups of single measurements in whole individual wells (e.g. mean intensity of a protein, E:I ratio, or edge strength). Wells with the same treatments from different plates were pooled together for comparisons, so treatment groups include equal contributions from 4 biological repeats. P-values are derived from 2-sided t-tests on Treatment and NonT sets of 11-18 wells.

UMAP dimensionality reduction and cluster analysis were performed on a combined random sample of 500 synapses within each well. Prior to UMAP analysis and Bayesian network inference, all protein measures were scaled to have a standard deviation of 1.

#### Bayesian network analysis and controlled edge calculation

Bayesian network inference was performed on a combined random sample of synapses from each well, while limiting only to excitatory-labeled synapses and 8 excitatory synaptic proteins. Measurements in each protein were discretized into 51 bins in the following way: all measurements of 0 (no puncta of that protein associated with the synapse) were assigned to bin 0, and bins 1-50 were assigned by equal-frequency discretization.

The discretized dataset was then sampled for 3000 points which were used to construct a Bayesian network using the likelihood-score-maximizing ‘tabu’ algorithm^8^. 50 such samplings and rederivations of the network were used to establish confidence in the presence and direction of edges. Network derivation was done using the tabu and boot.strength functions in the R package bnlearn^8^. A similar procedure was applied to simulated datasets, data from a previous synaptic scaling study^4^, and adversarially modified data sets.

Given a network, we define the strength of an edge between two nodes as the average correlation of the two variables across strata where the other parents of the daughter node are held constant^9,10^. That is, the strength of an edge from A to B, where B also has edges leading to it from C and D, as the correlation between A and B when controlling for C and D. To estimate that, we repeated the following algorithm for average correlations between A and B across strata of equal C and D:

- Sample a point (A_0_, B_0_, C_0_, D_0_)
- Find set of all points (A, B, C, D) such that 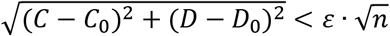 where n is the number of variables to control for (2 in this example) and ε is a predetermined tolerance level set at 0.5 (smaller tolerances did not yield significantly different measures)
- If the set contains more than 5 points, calculate Pearson’s correlation coefficient *cor*(*A*, *B*) across that set.
- Average the resulting correlation measure across 20 · 2^*n*^ such samplings.

A similar stratification procedure was done to assess the conditional effect of a certain treatment on protein A when controlling for proteins B and C. The treatment and NonT groups were pooled together, a point was sampled at random and a set of all points with similar B and C was found, and the log_2_-fold difference between the mean levels of A in treated vs NonT synapses was calculated and averaged across many samplings.

### Supplementary Tables

**Table S1:** Conjugated antibodies used in this study and additional antibodies for validation

| Target | Supplier and Cat. No. | Host, clonality and isotype | Imaged with |
| --- | --- | --- | --- |
| MAP2 | Novus Biologicals NB300-213 | Chicken polyclonal | Secondary IF |
| PSD95 | Cell Signaling Technology 3450 | Rabbit monoclonal | Secondary PRISM |
| Gephyrin | Synaptic Systems 147208 | Rat monoclonal IgG1 | Secondary PRISM |
| GluR2 | Synaptic Systems 182 105 | Guinea Pig polyclonal | Secondary PRISM |
| NR2A | Neuromab 75-288 | Mouse monoclonal IgG2a | Secondary PRISM |
| Shank3 | Synaptic Systems 162311 | Mouse monoclonal IgG1 | Secondary PRISM |
| anti-Rabbit | Invitrogen A16126 | Goat polyclonal | PRISM – p1 |
| anti-Mouse IgG1 | Abcam ab98689 | Goat polyclonal | PRISM – p4 |
| anti-Guinea Pig | Invitrogen A18777 | Goat polyclonal | PRISM – p6 |
| anti-Rat | Invitrogen A18873 | Goat polyclonal | PRISM – p7 |
| anti-Mouse IgG2 | Novus Biologicals nb7513 | Goat polyclonal | PRISM – p12 |
| F-actin | BACHEM H7643 | Phalloidin | Direct PRISM – p2 |
| vGAT | Synaptic Systems 131011 | Mouse monoclonal IgG3 | Direct PRISM – p3 |
| Bassoon | Enzo Life Sciences ADI-VAM-PS003 | Mouse monoclonal IgG2a | Direct PRISM – p8 |
| Synapsin1 | Synaptic Systems 106011 | Mouse monoclonal IgG1 | Direct PRISM – p9 |
| Homer1 | Synaptic Systems 160011 | Mouse monoclonal IgG1 | Direct PRISM – p10 |
| vGlut1 | Synaptic Systems N1602 | Camelid sdAb | Direct IF |
| F-actin | Abcam ab176759 | Phalloidin | Direct IF |
| mGluR5 | Sigma Aldrich AB5675 | Rabbit polyclonal | Secondary IF |
| Shank1/2/3 | Synaptic Systems 162111 | Mouse monoclonal IgG2a | Secondary IF |
| NR1 | Millipore MAB363 | Mouse monoclonal IgG2a | Secondary IF |
| anti-Mouse 647 | Abcam ab150107 | Donkey polyclonal |  |
| anti-Rabbit 568 | Abcam ab175470 | Donkey polyclonal |  |

**Table S2:**
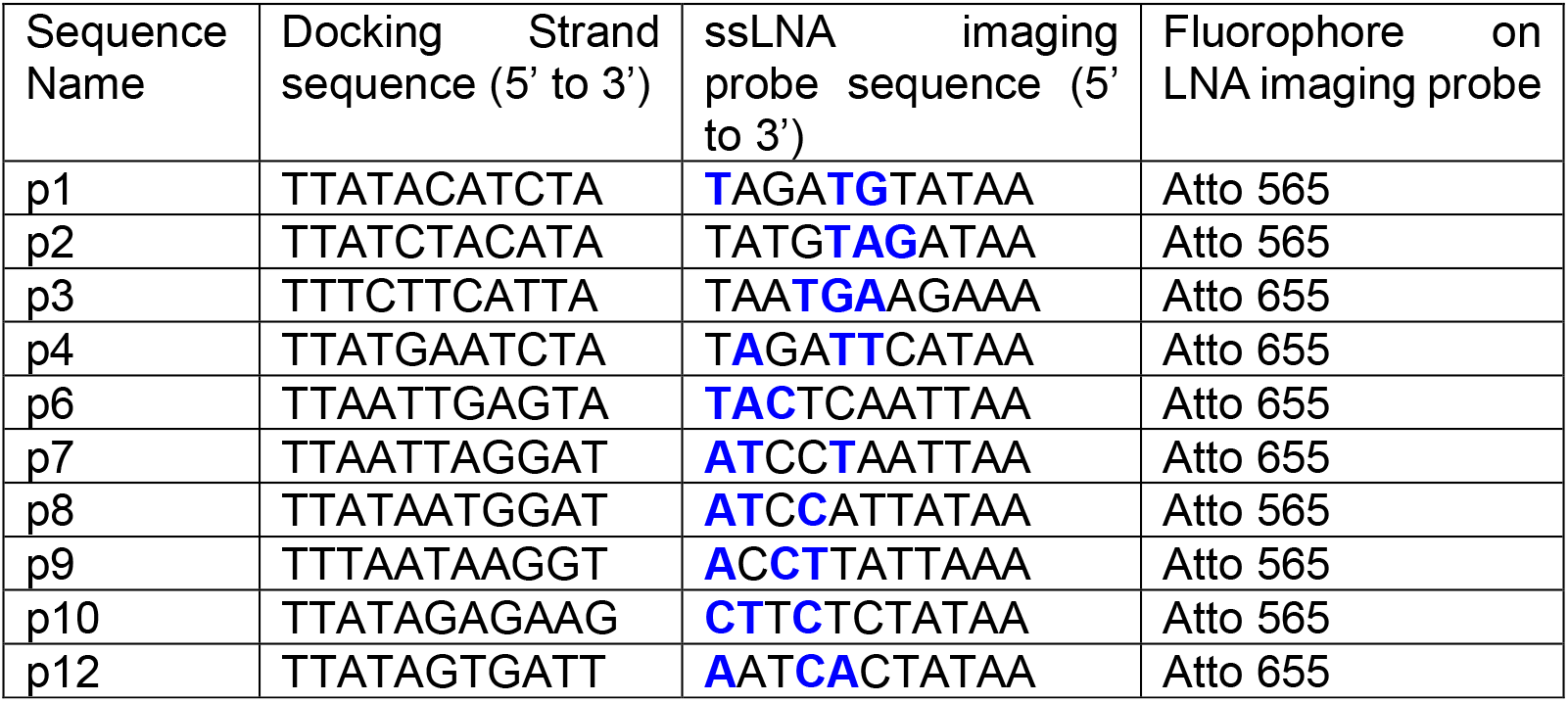
Docking and Imaging strand sequences:

**Table S3:** Accell siRNA mixtures and TaqMan RTqPCR primers:

| Gene | Accell siRNA Cat. No. | TaqMan RTqPCR Assay ID |
| --- | --- | --- |
| *Nontargeting* | D-001910-10-20 |  |
| *Dyrk1a* | E-090285-00 | Rn00562940_m1 |
| *Trio* | E-087912-00 | Rn01524285_m1 |
| *Cul1* | E-094875-00 | Rn01501284_m1 |
| *Setd5* | E-095571-00 | Rn01425934_m1 |
| *Sp4* | E-090979-00 | Rn00562717_m1 |
| *Chd8* | E-098578-00 | Rn00576005_m1 |
| *Herc1* | E-101129-00 | Rn01421830_m1 |
| *Shank3* | E-080173-00 | Rn00572344_m1 |
| *Scn2a* | E-097072-00 | Rn00680558_m1 |
| *Gria3* | E-088270-00 | Rn00583547_m1 |
| *Cacna1g* | E-089308-00 | Rn00709287_m1 |
| *Fmr1* | E-091941-00 | Rn00709627_m1 |
| *Syngap1* | E-080169-00 | Rn00710435_m1 |
| *Setd1a* | E-051358-00 | Mm00626141_m1 |
| *Grin2a* | E-091573-00 | Rn00561341_m1 |
| *Pten* | E-080104-00 | Rn00477208_m1 |
| *Xpo7* | E-087157-00 | Rn01493200_m1 |
| *Actb* |  | Rn00667869_m1 |

### Supplementary Figures

**Figure S1:**
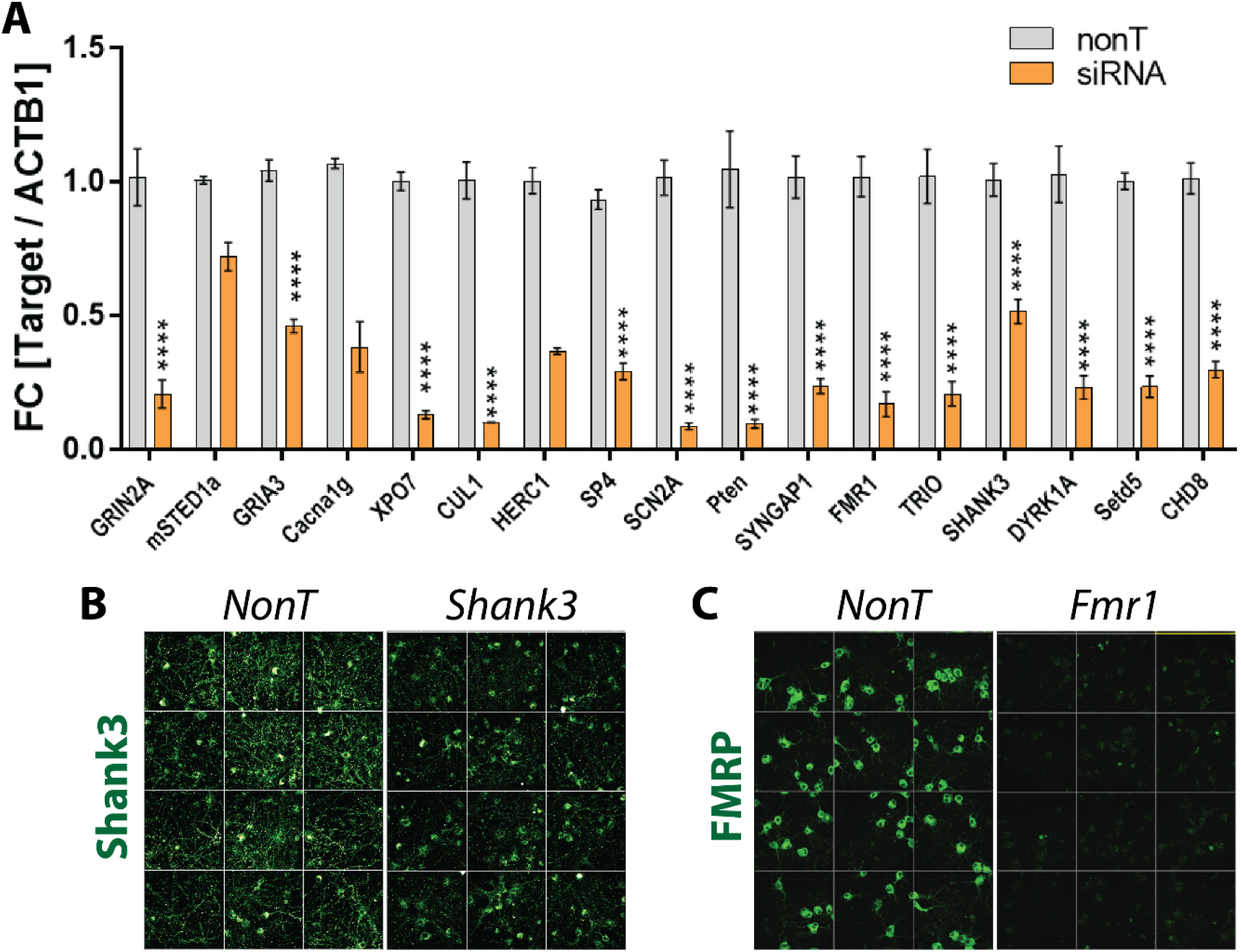
RNAi knockdown validation. A) RTqPCR of transcript abundances of the perturbed genes in rat embryonic hippocampal neurons after siRNA treatment at DIV 5 and harvest at DIV 21. Y-axis is normalized to NonT and to ACTB1 mRNA levels. Error bars indicate SEM across cultures. ****p<0.001. B,C) Immunofluorescence of Shank3 (B) and FMRP (C) in siRNA-treated vs NonT-treated cultures.

**Figure S2:**
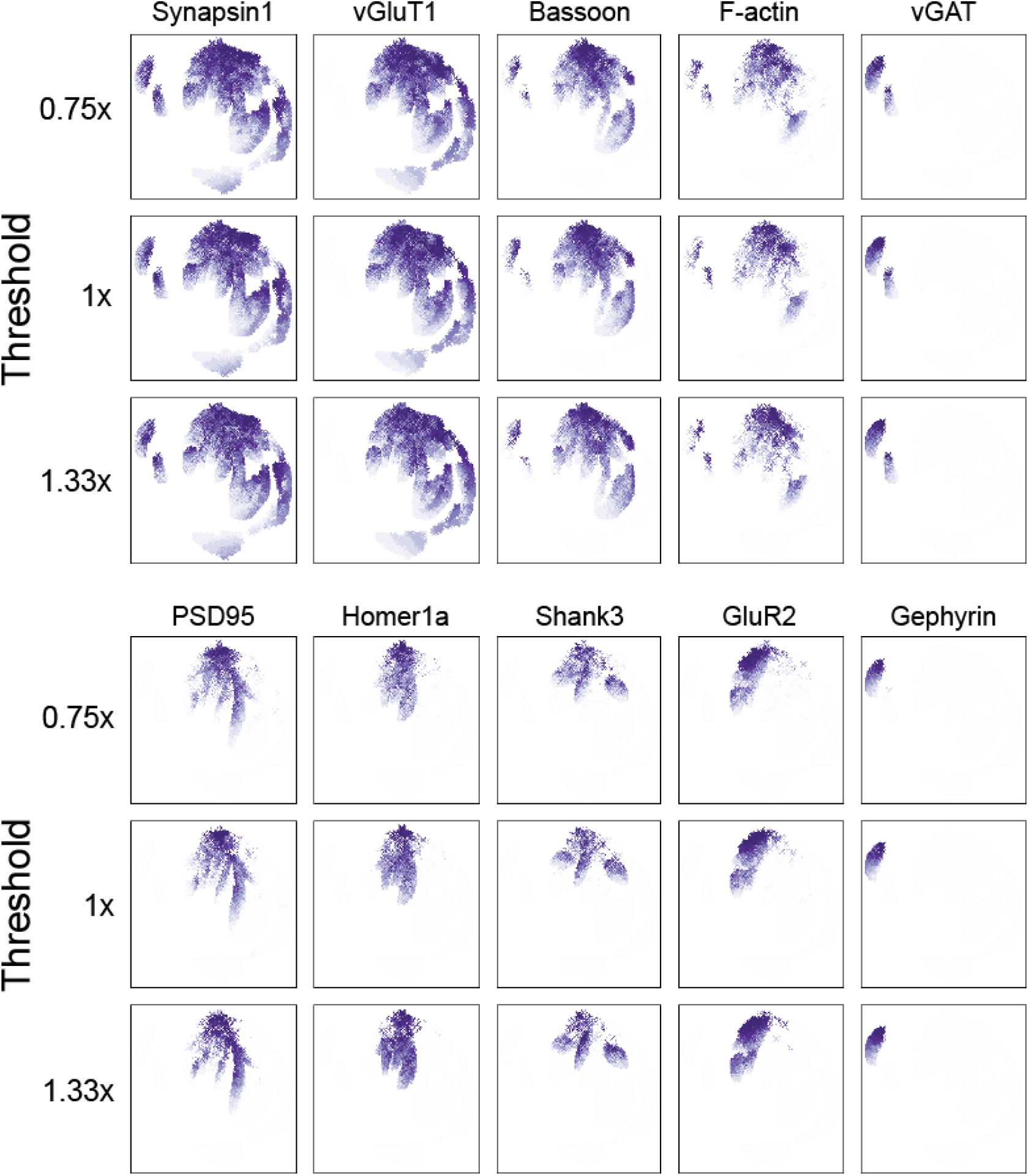
UMAP plot of data sampled from synapses thresholded at three levels, colored by protein level. Synapses for each threshold exhibit clusters defined by presence and absence of protein combinations.

**Figure S3:**
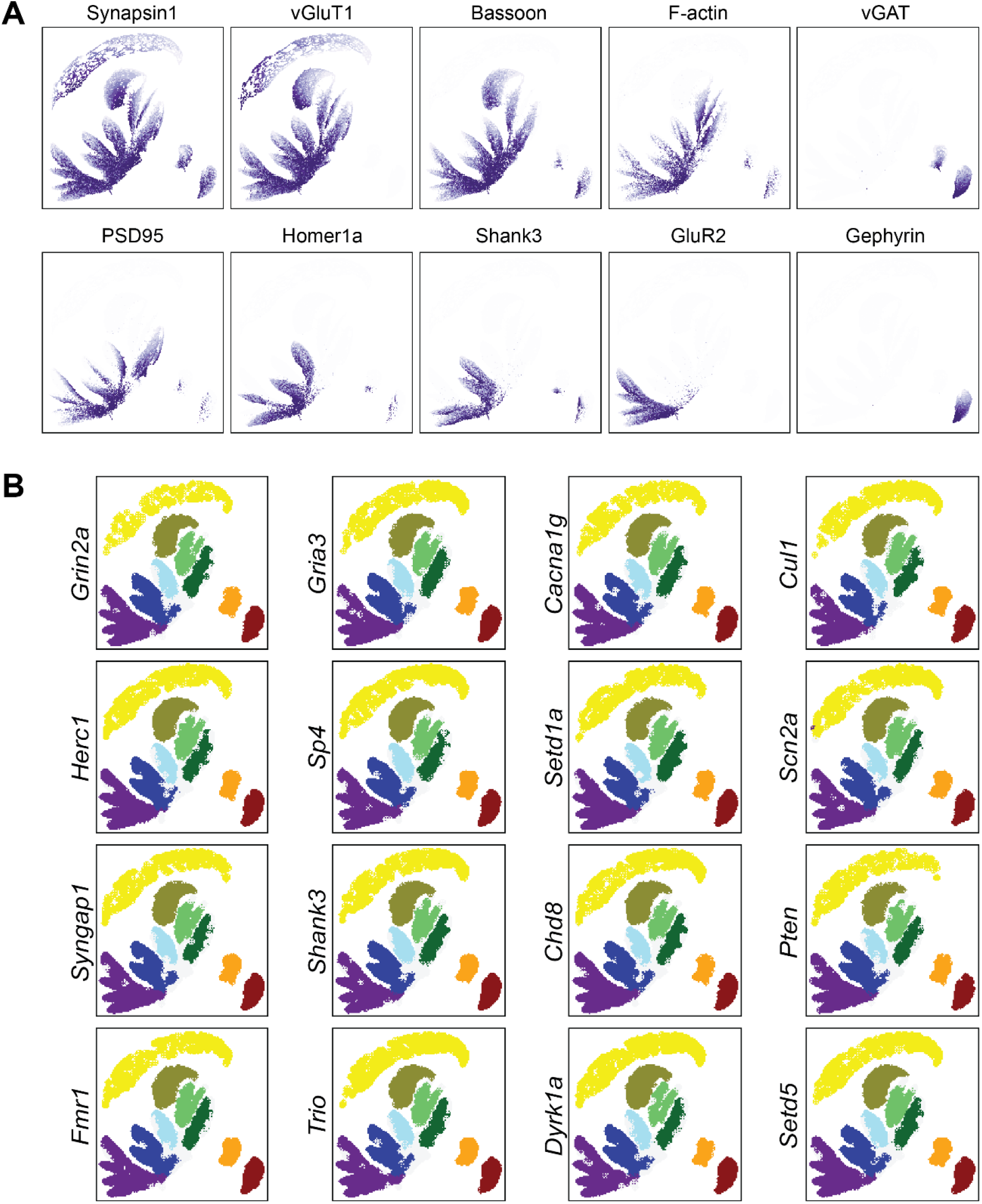
A) UMAP projections colored by levels of individual proteins. B) UMAP projections of synapses from each treatment group.

**Figure S4:**
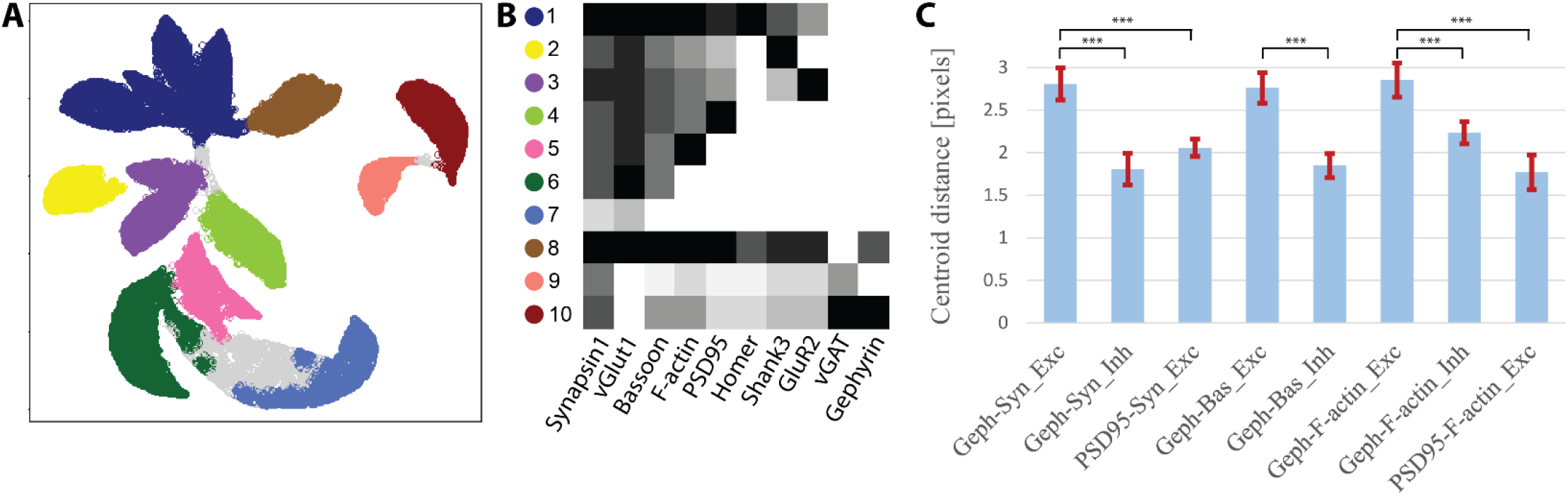
A,B) UMAP projection and unsupervised density-based clustering (A) of all synapses, and heatmap of average protein levels per cluster (B) similar to figure 3 but with an additional cluster (#8) of synapses positive for Gephyrin and vGluT. C) Average distances (in pixels) between centroids of puncta of different proteins in synaptic subsets. Left to right: Gephyrin-Synapsin in cluster #8, Gephyrin-Synapsin in cluster #10, PSD95-Synapsin across all PSD95+ excitatory synapses, Gephyrin-Bassoon in cluster #8, Gephyrin-Bassoon in cluster #10, Gephyrin-F-actin in cluster #8, Gephyrin-F-actin in cluster #10, and PSD95-F-actin across all PSD95+, F-actin+ excitatory synapses. Error bars are standard deviations across all wells. ***p<0.0001

**Figure S5:**
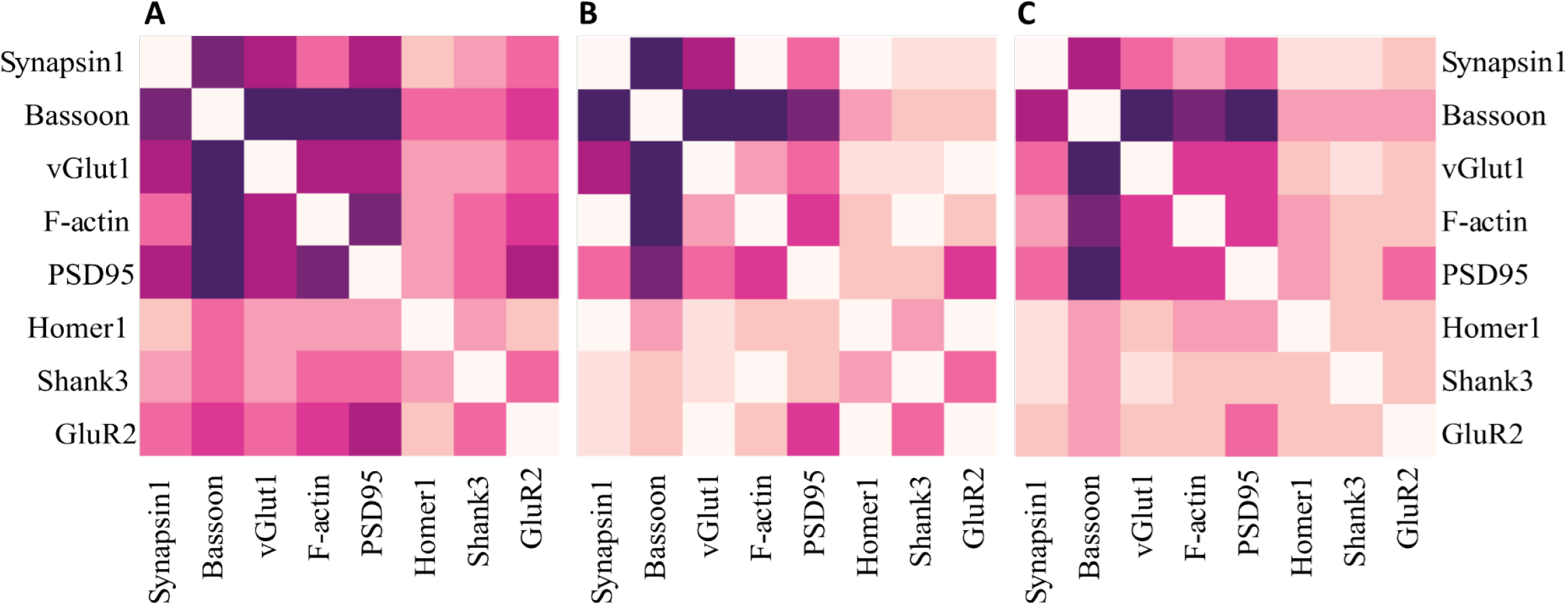
A) Direct correlations between proteins in excitatory synapses. B) Correlations in each pair, controlling for all other 6 proteins. C) Mutual information between protein levels

**Figure S6:**
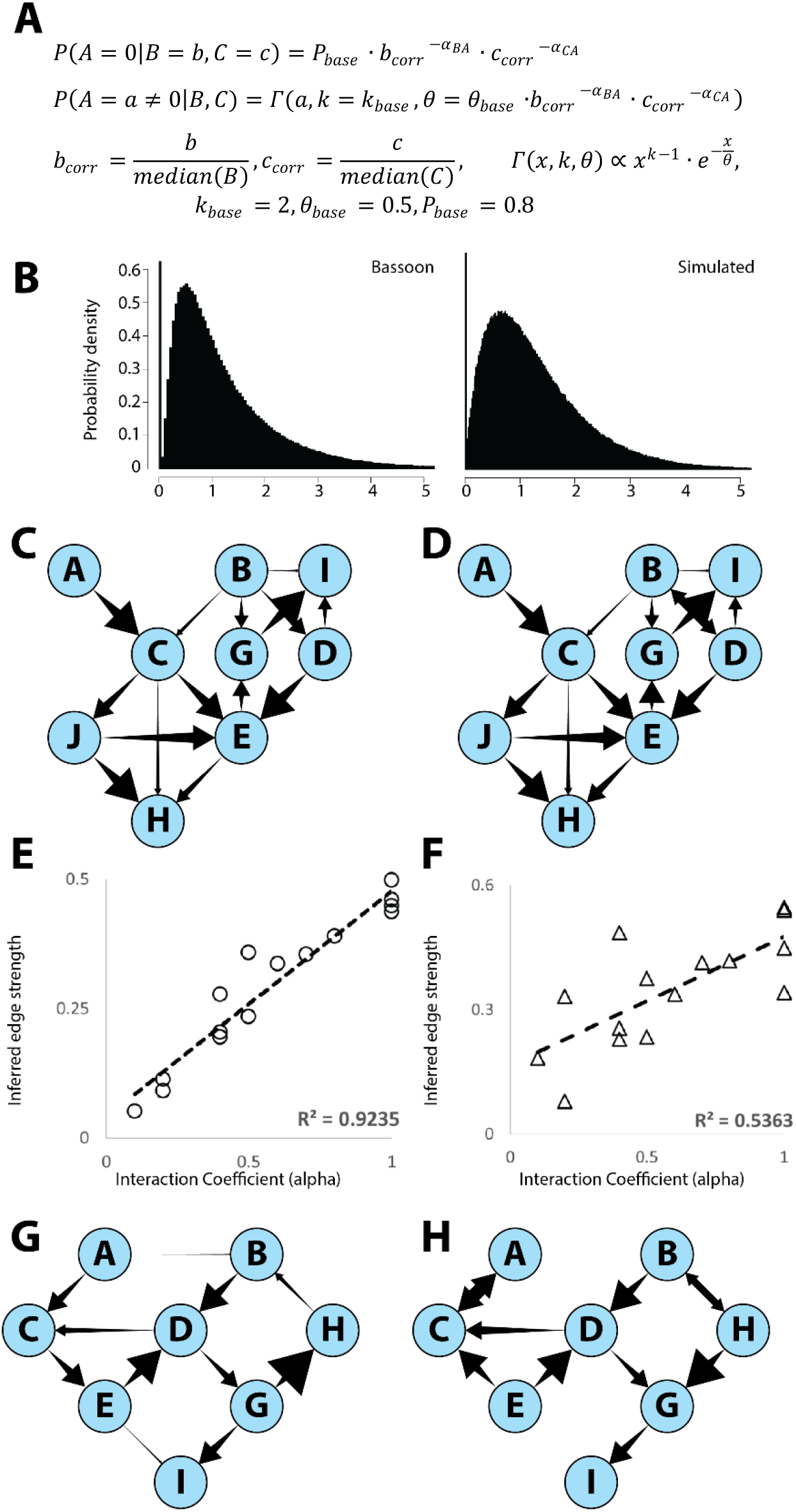
Bayesian network inference on simulated networks. A) Formulae to sample the simulated conditional distributions. B) Histograms of Bassoon values from excitatory PRISM data (left) and a simulated distribution with an equal proportion of zero values and equal mean. C) Simulated Bayesian network. D) Reconstructed network. E,F) Calculated edge strengths of the edges in (D) versus defined interaction coefficients in (C). E – edge strengths calculated as in figure 4A, by parent-controlled correlations. F – total (uncorrected) correlations. G) Simulated non-Bayesian network with cycles (CED and BDGH). Cycles were created by adding D→ C_2_ and H→B_2_ edges post-factum and defining C_final_=average(C,C_2_), B_final_=average(B,B_2_). H) Reconstructed Bayesian network. Some sensitivity to very weak interactions is lost and a few edge directions are scrambled but the overall structure and relative edge strengths are preserved.

**Figure S7:**
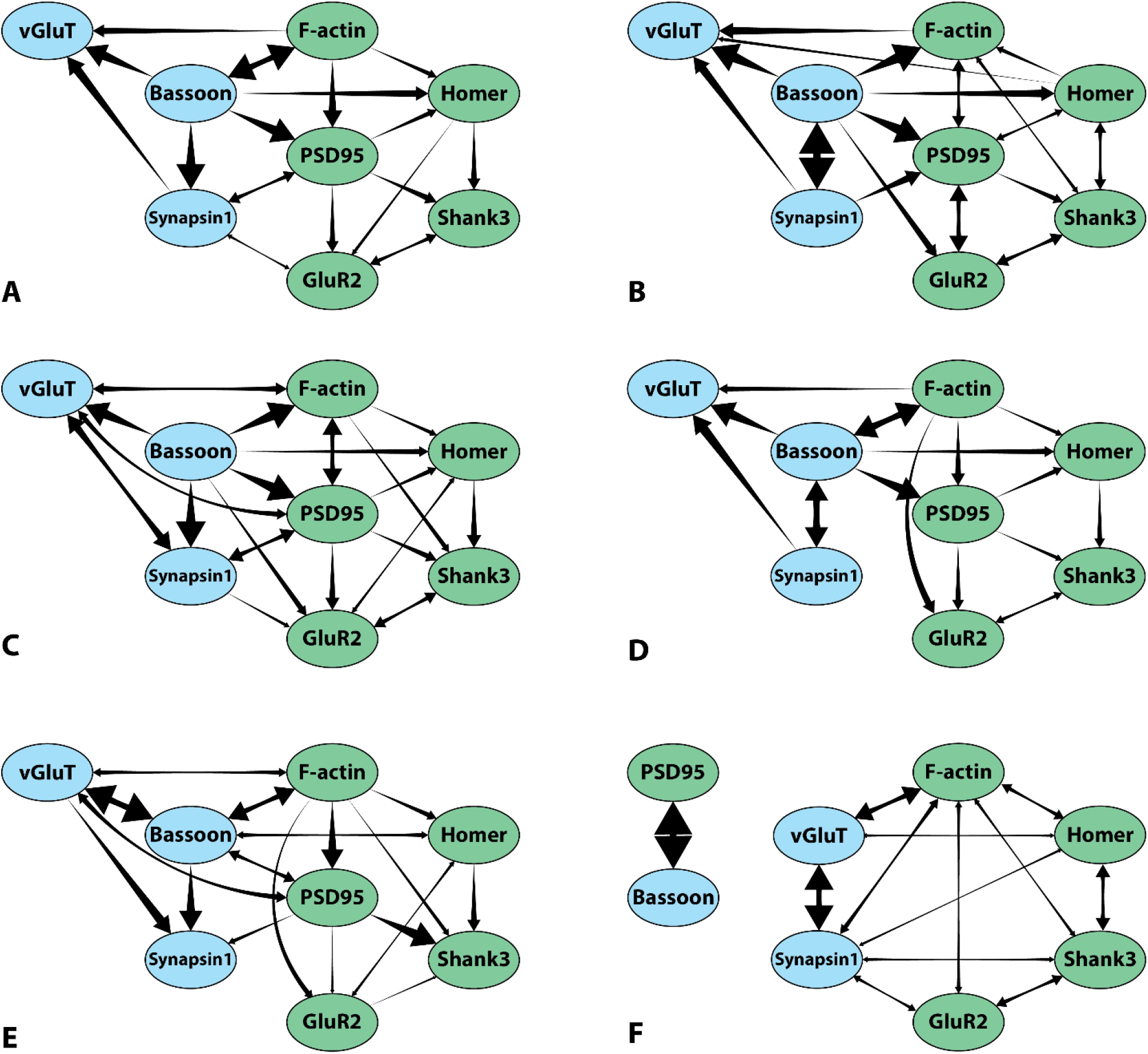
Bayesian networks derived from adversarially modified data. A) Original network. B) Network derived only from the subset of synapses which are positive for all components. C) Network derived on data with 25% lower threshold. D) Network derived on data with 33% higher threshold. E) Network derived on data in which synapses were identified (during CellProfiler analysis) not by Synapsin1 puncta but by a combination of F-actin and PSD95 puncta. F) Network derived from the original data, but with data for Bassoon and PSD95 levels randomly permuted.

**Figure S8:**
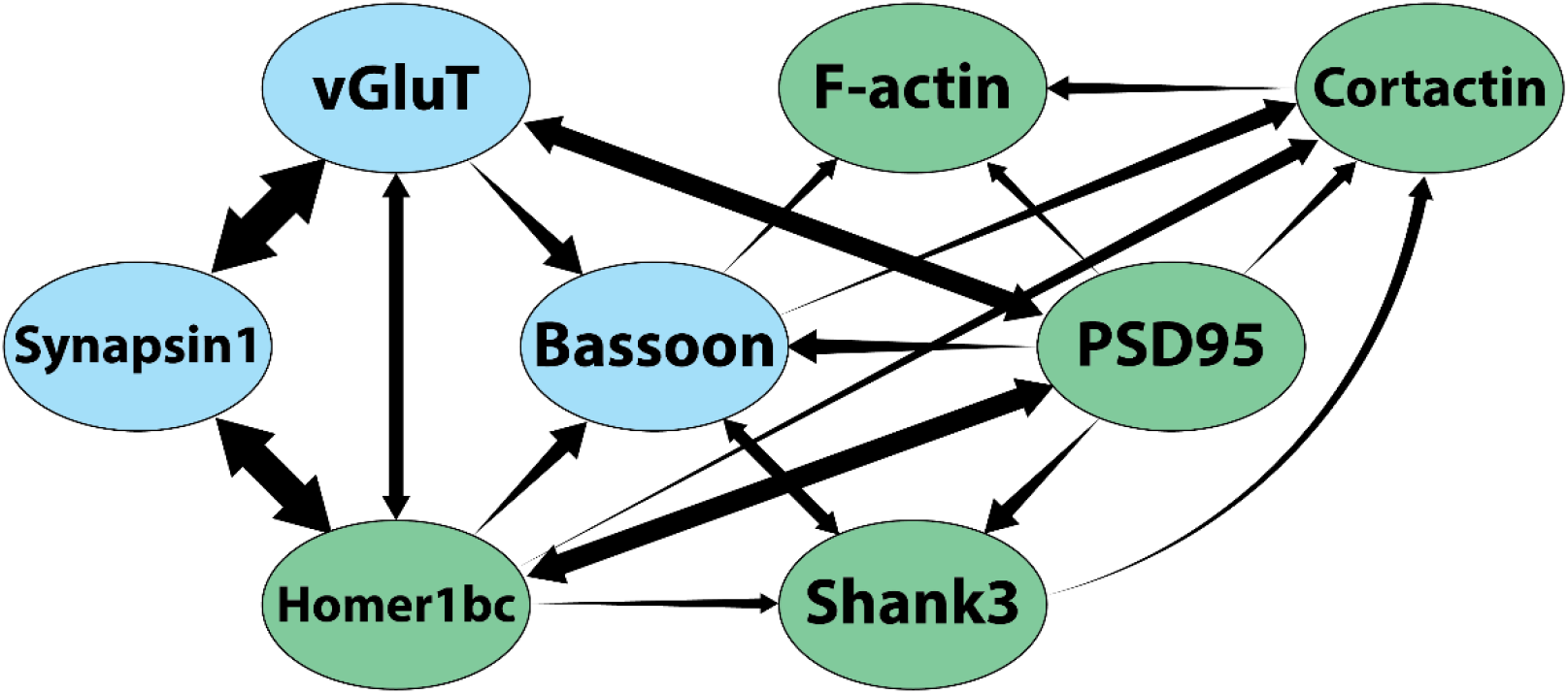
Bayesian network derived on data from a previous study. 6 proteins are the same in both studies – Synapsin1, vGluT1, Bassoon, Shank3, F-actin and PSD95 – but different antibodies were used. Homer1bc was replaced by Homer1a in this study, and Cortactin was replaced by GluR2.

**Figure S9:**
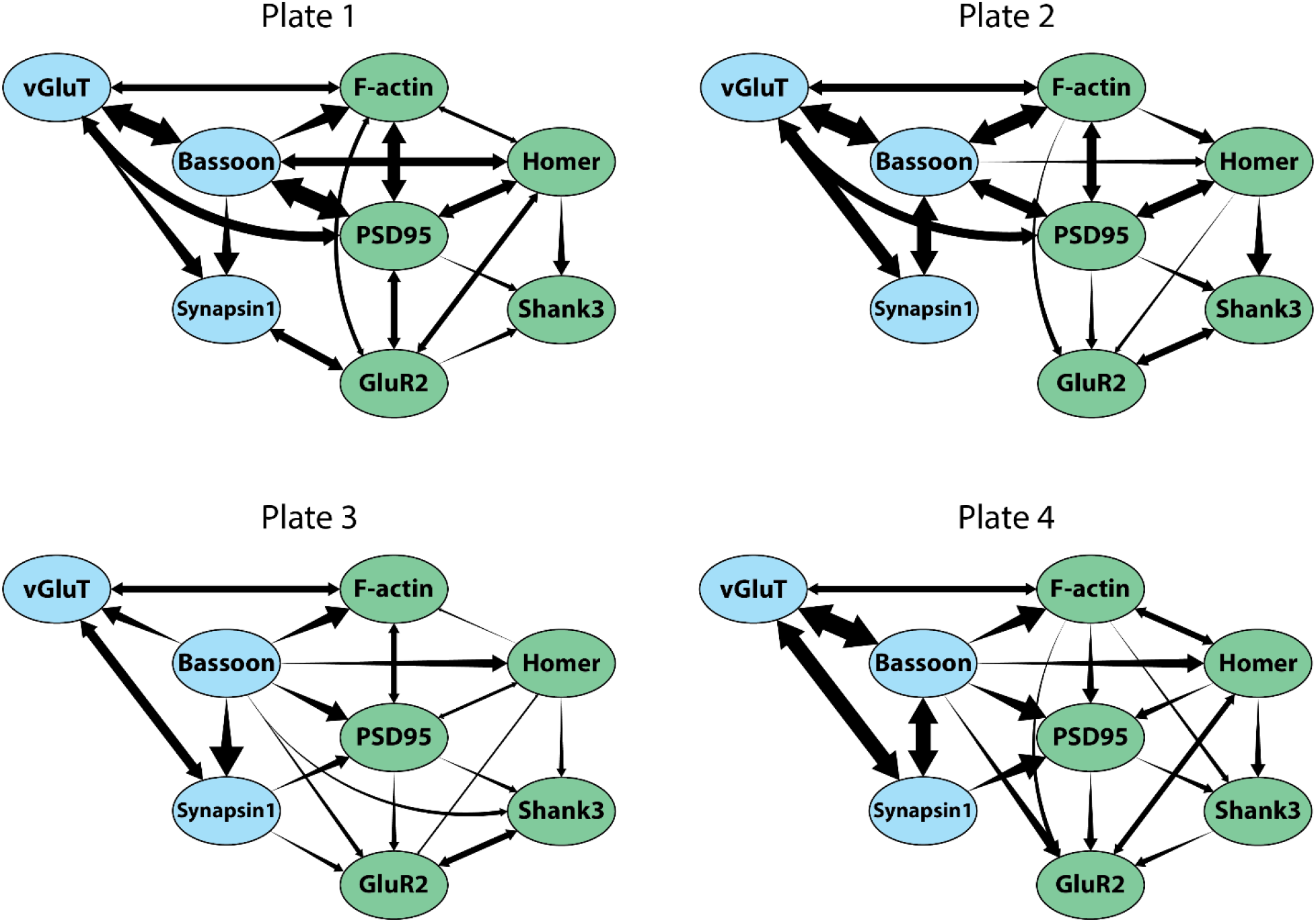
Bayesian networks derived on data from each experiment separately.

**Figure S10:**
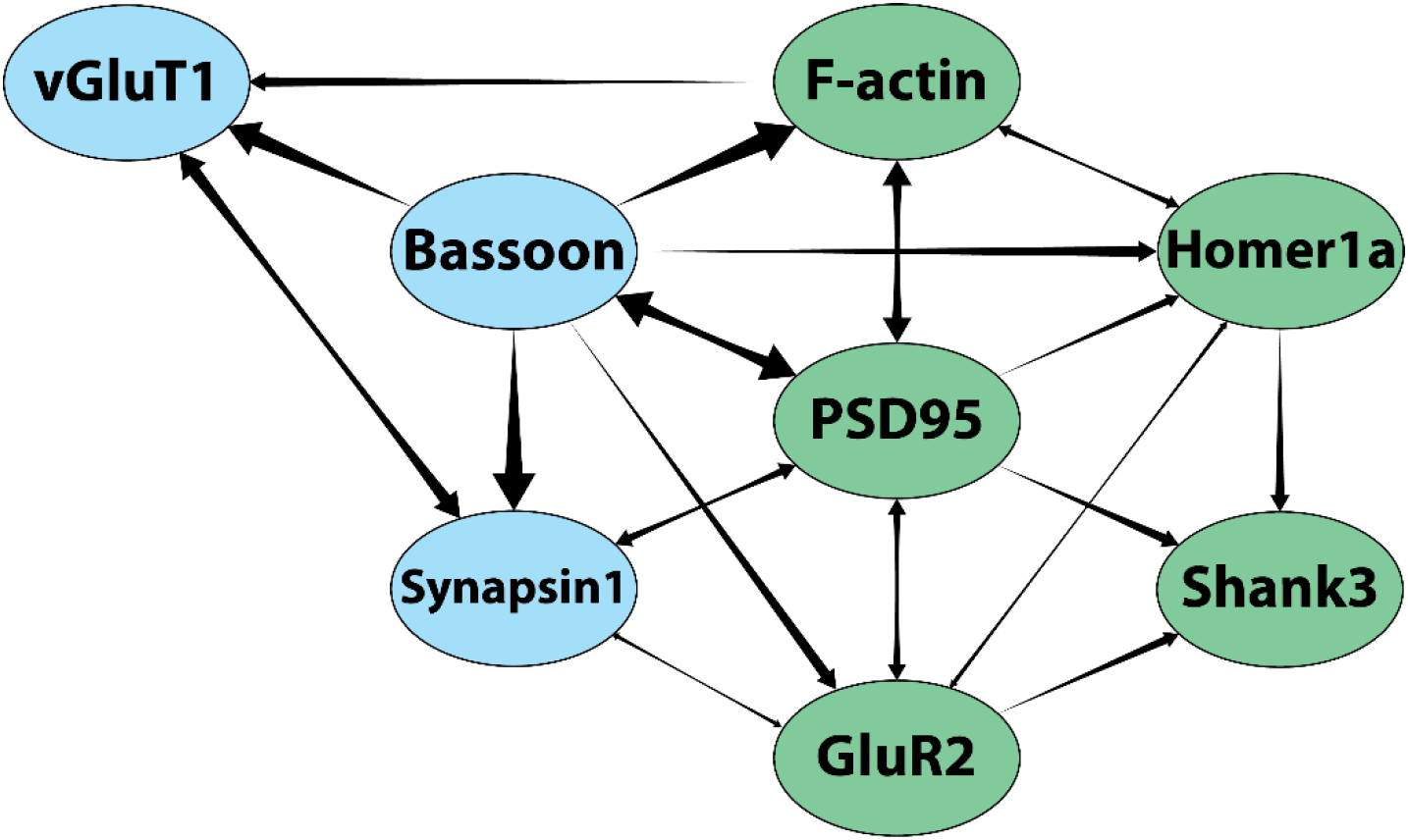
Bayesian network derived on data with *Shank3* knockdown cultures excluded

## Notes

### Competing Interest Statement

The authors have declared no competing interest.

